# Rate-limiting transport of positive charges through the Sec-machinery is integral to the mechanism of protein transport

**DOI:** 10.1101/592543

**Authors:** William J. Allen, Robin A. Corey, Daniel W. Watkins, A. Sofia F. Oliveira, Kiel Hards, Gregory M. Cook, Ian Collinson

**Affiliations:** School of Biochemistry, University of Bristol, University Walk, Bristol BS8 1TD, UK; School of Chemistry, University of Bristol, University Walk, Bristol BS8 1TD, UK; Department of Microbiology and Immunology, University of Otago, Dunedin 9054, New Zealand

## Abstract

Transport of proteins across and into membranes is a fundamental biological process with the vast majority being conducted by the ubiquitous Sec machinery. In bacteria, this is usually achieved when the SecY-complex engages the cytosolic ATPase SecA (secretion) or translating ribosomes (insertion). Great strides have been made towards understanding the mechanism of protein translocation. Yet, important questions remain – notably, the nature of the individual steps that constitute transport, and how the proton-motive force (PMF) across the plasma membrane contributes. Here, we apply a recently developed high-resolution protein transport assay to explore these questions. We find that pre-protein transport is limited primarily by the diffusion of arginine residues across the membrane, particularly in the context of bulky hydrophobic sequences. This specific effect of arginine, caused by its positive charge, is mitigated for lysine which can be deprotonated and transported across the membrane in its neutral form. These observations have interesting implications for the mechanism of protein secretion, suggesting a simple mechanism by which PMF can aid transport, and enabling a ‘proton ratchet’, wherein re-protonation of exiting lysine residues prevents channel re-entry, biasing transport in the outward direction.

## Introduction

The final destination of proteins synthesised in the cytosol is governed by cleavable N-terminal signal sequences (SS), necessary and sufficient for targeting (Blobel and Dobberstein, 1975), or by information in the mature protein. They are either retained and subject to controlled folding, or are recognised by factors for delivery, usually in an unfolded state, to bespoke protein translocation machineries (translocons) for carriage across or into cellular membranes. Amongst these, and responsible for the majority both of protein secretion and membrane protein insertion, is the universally conserved Sec system: at its core the heterotrimer SecYEG/β in plasma membrane of bacteria and Archaea or Sec61αβγ in the endoplasmic reticulum of eukaryotes. The Sec system transports unfolded proteins, either immediately as they emerge from the ribosome (co-translationally) or post-translationally (Arkowitz et al., 1993).

In bacteria, membrane protein insertion is generally co-translational, whereas almost all secretion is post-translational, mediated by the cytosolic ATPase SecA (Lill et al., 1990). During passage across the membrane, pre-proteins with a SS are recognised by SecA, either directly or with the assistance of chaperones such as SecB, and brought to SecYEG in the membrane (Fig. 1; Hartl et al., 1990; Kumamoto and Beckwith, 1983; Oliver and Beckwith, 1982). The SS unlocks the translocon and initiates transport by inserting as a hairpin through the channel (step *i* in Fig. 1; Corey et al., 2016b; Ma et al., 2019), using ATP hydrolysis by SecA as an energy source (Economou and Wickner, 1994; Fessl et al., 2018; Lill et al., 1990). Transport of the rest of the of the polypeptide substrate proceeds stepwise (steps *ii-v*, elaborated below), with cycles of ATP turnover (Brundage et al., 1990), aided *in vivo* by the electrochemical gradient of protons across the membrane – the proton-motive force (PMF; positive and acidic outside; (Schiebel et al., 1991). Once the pre-protein has crossed the membrane, the SS is cleaved off by signal peptidase and the transported protein released (Josefsson and Randall, 1981); this release step appears to require additional factors – most likely the periplasmic chaperone PpiD (Antonoaea et al., 2008; Fürst et al., 2018) – as it is very slow when measured *in vitro* using *ex vivo* or purified and reconstituted components (Allen et al., 2020; Mao et al., 2020).

**Figure 1.**
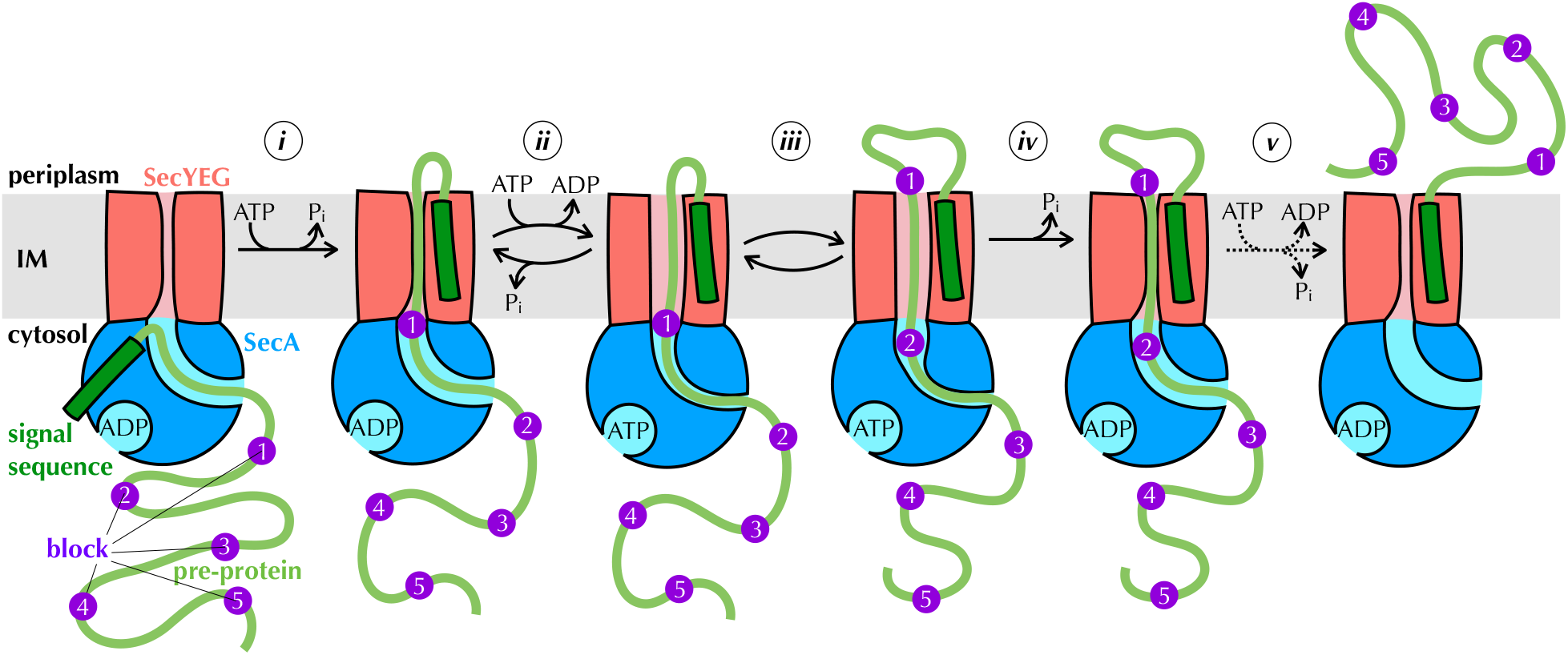
The proton ratchet model. Brownian ratchet model of pre-protein transport, based on (Allen et al., 2016). Cycles of ATP binding and hydrolysis by SecA (blue) allow blocks (purple) in the pre-protein substrate (green) to diffuse outwards through SecYEG (red) one at a time, but backsliding is prevented, leading to directional transport. See text for more detail.

Results from our laboratory and elsewhere, using the model *Escherichia coli* Sec system, have shown how ATP turnover could be coupled to directional pre-protein movement (Allen et al., 2016; Catipovic et al., 2019; Catipovic and Rapoport, 2020; Corey et al., 2019). Adding a non-hydrolysable ATP analogue to the system, which emulates the effect of nucleotide exchange (step *ii* in Fig. 1) has two major observable effects: it widens the channel through SecYEG (Allen et al., 2016) while tightening the clamp around pre-protein in SecA (Ahdash et al., 2019; Catipovic et al., 2019). Nucleotide exchange itself is promoted by perturbation of the two-helix finger (2HF; Allen et al., 2016; Zimmer et al., 2008) of SecA caused by the presence of pre-protein at the entrance to the channel through SecYEG (Allen et al., 2016). Together, these observations led us to propose a model in which pre-protein moves through the channel primarily by diffusion (Allen et al., 2016). Sequences that cannot diffuse across the membrane in the ADP-bound state (blocks, purple in Fig. 1) trigger nucleotide exchange (step *ii*), opening the SecYEG channel to allow them to slide through (step *iii*) and simultaneously clamping SecA shut to prevent them slipping backwards. ATP hydrolysis resets the channel (step *iv*), trapping blocks sequences that have diffused across on the outside of the membrane, thus providing directionality. This process (steps *ii-iv*) is repeated for each remaining block (step *v*) until the entire pre-protein has crossed the membrane.

A major prediction of the above model is that not every ATP turnover will give rise to a transport event, as step *ii* is reversible. Recent high resolution protein transport data, collected using a new assay based on split NanoLuc luciferase (Dixon et al., 2016; Pereira et al., 2019), lent support to this notion. In this assay, the large fragment of NanoLuc (11S) is encapsulated within proteoliposomes (PLs) or inverted membane vesicles (IMVs) containing SecYEG, while a high affinity small fragment (pep86) is incorporated into the pre-protein (Fig. 1 – Figure Supplement 1). As soon as pep86 enters the vesicle it combines with 11S, producing a luminescence signal (Fig. 1 – Figure Supplement 1; Allen et al., 2020; Pereira et al., 2019)). Using detailed kinetic modelling, we showed that transport occurs in a relatively small number of steps – about five in the case of the model pre-protein pSpy – and that each step requires many ATP turnovers to resolve (an estimated 120 for pSpy *in vitro*, see Allen et al., 2020). However, we were unable to show exactly what these steps – presumably the blocks in the above model – physically correspond to. It also remains unclear how the PMF contributes.

To answer these questions, we have now generated a range of different pSpy variants with different physical and chemical properties. Employing the NanoLuc assay to measure their transport both in the presence and absence of proton-motive force (PMF), we show that transporting positively charged residues across the membrane is the slowest step of transport, with bulkier residues also contributing. This strong barrier to positive charges is partially overcome by deprotonating lysines at the cytosolic face of the membrane; for this reason, arginines, which have a much higher p*K*_a_ than lysine, are by far the hardest amino acid to transport. Surprisingly, however, the PMF does not appear to exploit the net negative charge this confers on the polypeptide to move it through the channel electrophoretically; rather, charged residues seem unaffected by the electrical component of PMF (Δψ) in our experimental setup.

Our results provide important new mechanistic insights of the mechanism of protein secretion as the step size can help distinguish between alternative models for protein transport. A diffusional based system described above will have variable steps and step sizes depending on the nature of the polypeptide. This is because the forward biasing of diffusional transport will be affected by the ratcheting potential of the translocating polypeptide. In contrast, a power-stroke mechanism relying on an ATP-dependent piston movement of SecA that pushes a stretch of polypeptide across the membrane (Catipovic et al., 2019; Catipovic and Rapoport, 2020) will have an identical number of steps, irrespective of the nature of the polypeptide. Thus, variations in these parameters could prove decisive in distinguishing these mechanisms. In addition to the implications for the mechanism of protein secretion, the results also hint at the underlying basis for the ‘positive-inside rule’ for the topology of membrane proteins (Cymer et al., 2015; vonHeijne, 1989).

## Results

### The chemical and physical properties of pre-protein affect its transport characteristics

We started by investigating which physical properties of a pre-protein determine how fast it is transported through SecYEG, by systematically varying the amino acid sequence of the model Sec substrate pSpy. It quickly became apparent that altering pSpy affects other properties of the pre-protein in undesirable ways, including solubility, affinity for SecA, and possibly the rates of initiation and termination. Pre-proteins need to be presented to the assay in an unfolded state, which we do *in vitro* in 6 M urea solution; the protein substrate is then diluted out of urea into the assay mixture. If the solubility is perturbed then this results in precipitation of the substrate at this stage, affecting the result.

Therefore, to avoid detrimental changes of pre-protein behaviour, we created constructs consisting of the pSpy SS followed by three tandem mature (m)Spys, with a pep86 sequence after the second (pSpy_XLX_; XLX in Fig. 2a) – and altered only the central one. The two flanking native Spy sequences ensure that the beginning and end of transport are always the same, and prevent the less stable Spy variants from precipitating upon dilution out of urea. As a control, we created the same construct but with pep86 after the first mSpy (pSpy_LXX_, LXX in Fig. 2a). The difference in transport time between these two proteins (measured from the lag before transport signal appears, see Fig. 2b and Allen et al., 2020) corresponds exactly to the time it takes to transport the central mSpy (pink in Fig 2a,c). The lags for these match perfectly with those for the pSpy_4x_ series used previously (Allen et al., 2020; Fig. 2 – Figure Supplement 1a-b), confirming that lag is indeed a good, sensitive measure of transport rate.

**Figure 2.**
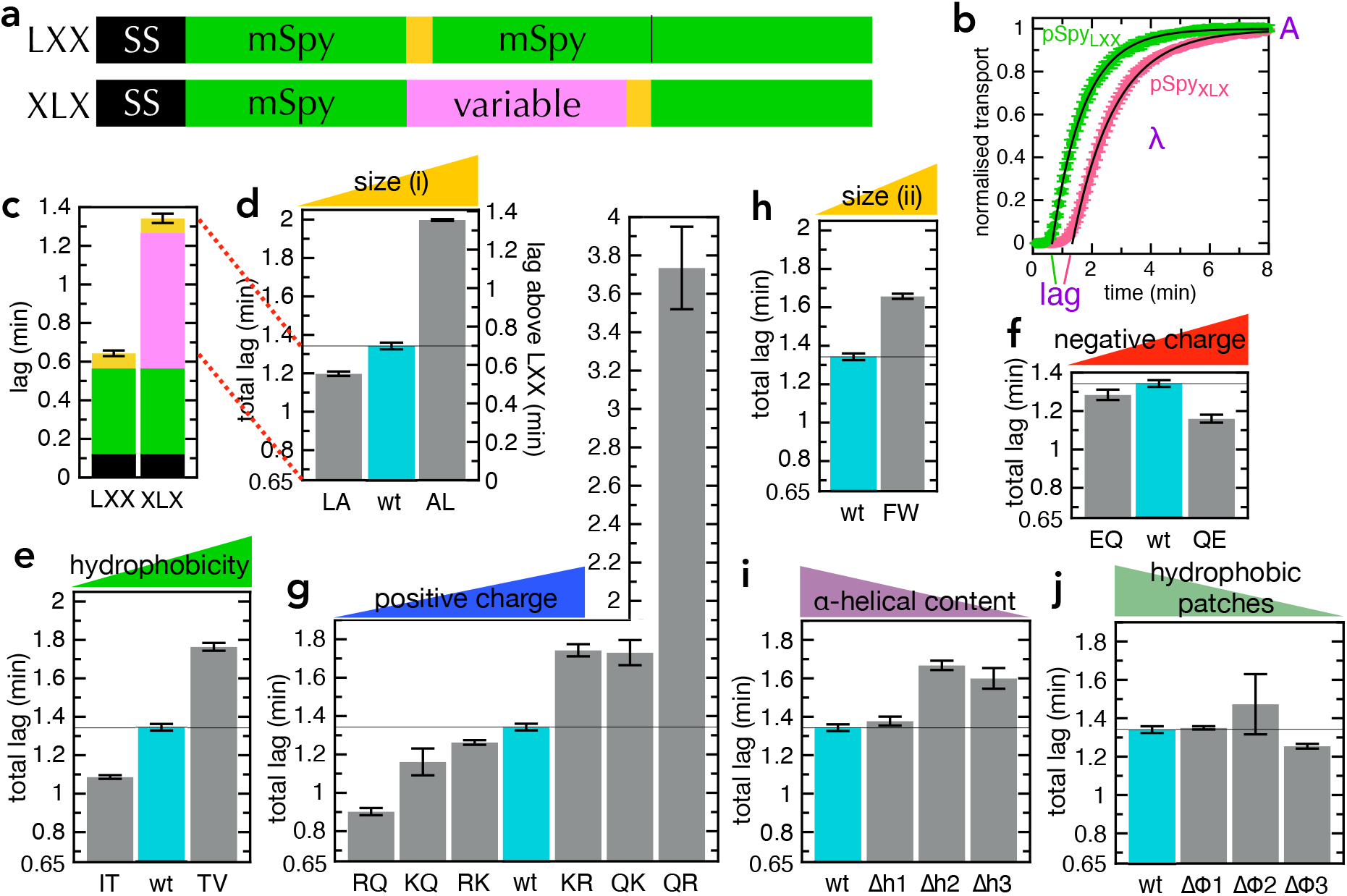
Transport of pSpy_XLX_ variants. **a)** Schematic of pSpy_LXX_ (LXX) and pSpy_XLX_ (XLX). Transport occurs from the N- to C-terminus (left to right), and a luminescence signal appears as soon as pep86 (yellow) enters the lumen of the PL. **b)** Transport traces of pSpy_LXX_ (green) and pSpy_XLX_(pink), normalised to give an amplitude (A; signal when the transport reaction reaches completion) of 1, and fitted to the simple lag + single exponential model (Allen et al., 2020). In this model, lag is the minimum time required for pre-protein transport, and corresponds to the sum of time constants for all transport steps (equal to 1/*k*, where *k* is the rate constant for that step). λ is a complicated variable, incorporating the transport rates but also the probability of transport pausing or failure and resetting (see Allen et al., 2020 for more details). Data are the average and SEM from 3 (pSpy_LXX_) or 11 (pSpy_XLX_) experimental replicates. **c)** Lag (taken from panel **b**) for pSpy_LXX_ and pSpy_XLX_. **d-j)** Lags for a range of pSpy_XLX_ variants, where the central mSpy has its chemical and physical properties varied (see text for details). In each case the y-axis starts at the transport time for pSpy_LXX_, so the visible part of the bar corresponds to the transport time only of the variable region. Data show the average and SEM from three experimental replicates.

Our first set of pSpy_XLX_ variants were created by replacing 6-8 of one residue with another, distributed as evenly as possible through mSpy (all sequences shown in Supplementary Table 1). We chose four general amino acid properties: size (L→A, A→L; Fig. 2d), hydrophobicity (T→V, I→T; Fig. 2e), negative charge (Q→E, E→Q; Fig. 2f), and positive charge (R→Q, Q→R, K→Q, Q→K, K→R, R→K; Fig. 2g). In the case of positive charge, we looked both at lysine – which can in some environments be deprotonated at physiological pH (Isom et al., 2011) – and arginine, which is generally considered to always retain its positive charge (Harms et al., 2011, and see also below). We find that transport is slower (longer lag than native pSpy_XLX_ above that of pSpy_LXX_, 0.65 min) when more bulky, hydrophobic and positively charged residues are present, while negative charges have limited effect. By far the strongest effect comes from changing the number of arginines: removing all eight (RQ in Fig. 2g) reduces the transport time three-fold (from 42 s to 16 s), while adding a further eight (QR in Fig. 2g) slows transport over four-fold (to 185 s).

Leucine is more hydrophobic than alanine, while threonine is somewhat smaller than isoleucine or valine (Fig. 2 – Figure supplement 2a), so these experiments do not distinguish particularly effectively between bulkiness and hydrophobicity. We therefore designed an additional variant in which all five phenylalanine residues were replaced with tryptophan (F→W; Fig. 2h). This increases bulkiness while decreasing hydrophobicity (Fig. 2 – Figure supplement 2a), allowing the two effects to be disentangled. The results reveal that the F→W substitutions slow transport, suggesting that residue size is a more important factor than hydrophobicity.

We have previously shown that the ATP hydrolytic cycle of SecA can influence the formation of secondary structure (primarily α-helix) within the translocating pre-protein (Corey et al., 2019). To explore the effect of this on transport kinetics, we designed pSpy_XLX_variants in which the helical propensity of one, two or three regions was reduced, without affecting the hydrophobic character of that region (Δh1, Δh2 and Δh3; Fig. 2i; Fig. 2 – Figure supplement 2b, purple). A converse set of mutations, in which the hydrophobic character is reduced without altering the helical propensity, was also generated (Δϕ1, Δϕ2 and Δϕ3; Fig. 2j; Fig. 2 – Figure supplement 2b, green). The transport parameters with these variants show that if sufficient helical content is removed it somewhat slows transport (Fig. 2i), supporting the notion that helix formation is part of the mechanism of transport (Corey et al., 2019). Removing hydrophobic patches, meanwhile, has marginal if any effect on transport rate (Fig. 2j), reaffirming that residue size is a bigger factor than hydrophobicity.

### Specific measurement of the transport steps for the pSpy_XLX_ variants

The lag is a useful measure of overall transport time, but it cannot distinguish between a large number (n) of fast steps (with rate *k*_step_) and a small number of slow steps. For this, we employed a numerical model of transport, implemented in Berkeley Madonna (Fig. 3 – Figure Supplement 1a; Allen et al., 2020, and see Methods). Just as described previously (Allen et al., 2020), best fits to the experimental data were calculated over a range of values for n, allowing *k*_step_ and *k*_fail_ to vary but fixing the other rate constants to previously determined values. Using this analysis, we find that transport of pSpy_LXX_ is best described by 5 steps with an average rate (*k*_step_) of 4.27 min^-1^ (green in Fig. 3a), while transport of pSpy_XLX_ proceeds in 9 steps with *k*_step_ = 4.19 min^-1^ (pink in Fig. 3a).

**Figure 3.**
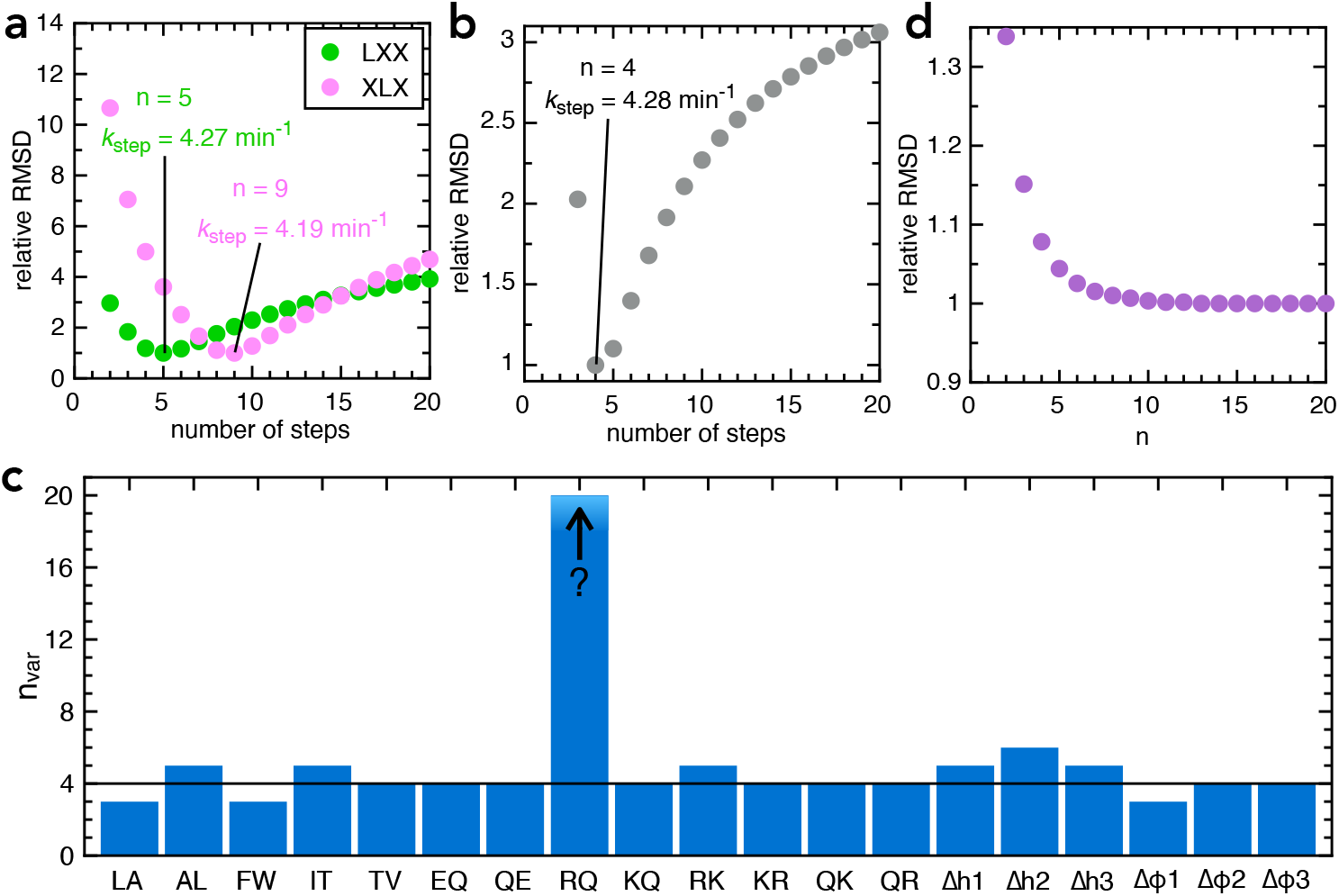
Determining step size for the variable mSpy regions. **a)** RMSD (normalised to its lowest value) for fits to the experimental transport data to the original transport model (see Figure 3 – Figure Supplement 1a and Allen et al., 2020), over a range of values for n (number of steps), using Berkeley Madonna. Values for other parameters are *k*_block_ = 0.31 min^-1^ (determined previously; Allen et al., 2020), *k*_on_ = 0.96 µM^-1^.min^-1^ (see Fig. 3 – Figure Supplement 2a) and k_off_ = 0.085 min^-1^ (see Fig. 3 – Figure Supplement 2b). Best fits are to 5 steps for pSpy_LXX_(green; *k*_step_ = 4.27 min^-1^) and 9 steps for pSpy_XLX_ (pink; *k*_step_= 4.19 min^-1^). **b)** RMSD (normalised to its lowest value) for fits of experimental transport data for pSpy_XLX_ to the model in Figure 3 – Figure Supplement 1b, over a range of values for n_var_. All parameters other than *k*_step,var_ and *k*_fail,var_ are fixed to the same values as in panel **a**. The best fit is to n_var_ = 4, *k*_step,var_= 4.28 min^-1^. **c)** Best fit n_var_ for each of the pSpy_XLX_ variants in Fig. 2d-j, calculated as in panel **b**, but with brightness adjusted for the values in Figure 3 – Figure Supplement 3 (see text for details), and *k*_block,var_ allowed to float. **d)** Normalised RMSD as a function of n_var_ for pSpy^R→^ (all parameters as in panel **c**).

To extract n specifically for transport of the second mSpy of pSpy_XLX_ (Fig. 2a, pink; the variable one in the above constructs; n_var_), we split the Berkeley Madonna model into two parts (Fig. 3 – Figure Supplement 1b). All parameters for transport of the first mSpy (including initiation and pep86 transport) were fixed to the best fit for pSpy_LXX_ (n = 5, *k*_step_= 4.27 min^-1^), while *k*_step_ for the variant (*k*_step,var_) was allowed to float and the best fit determined over a range of values for n_var_. This gives a best fit of n = 4 for pSpy_XLX_ (Fig. 3b), just as expected from the fit to the simpler model (Fig. 3a).

We next used this model to extract n_var_ for each of the 19 pSpy_XLX_ variants. The amplitudes of each variant (relative to native) differ somewhat from sequence to sequence (Fig. 2 – figure supplement 2, grey bars) – due either to differences in the signal produced by each NanoLuc (‘brightness’ in the Berkeley Madonna model) or in the probability that the sequences become trapped within the channel (*k*_block_). To distinguish these possibilities, we measured the NanoLuc signal of each variant in solution, under saturating conditions (Fig. 3 – Figure Supplement 2, turquoise bars). The results suggest that in most but not all cases, the variance is down to small differences in NanoLuc brightness (compare grey and turquoise bars in Fig. 3 – Figure Supplement 3). To account for this in the fitting we fixed brightness for each variant based on its measured value relative to native pSpy_XLX_, then allowed *k*_block,var_ to float, in addition to *k*_step,var_ and k_fail,var_.

The best fit number of steps for each variant is shown in Fig. 3c (full fitting results are in Supplementary Table 2). In all but one case, n falls between 3 and 6 (*i*.*e*. very similar to the native sequence, n = 4). Strikingly, however, it is not possible to determine a number of steps for pSpy ^R→Q^: the RMSD continues to go down then platueas as n_var_ increases (at least up to 20; Fig. 3d). We conclude that in the vast majority of cases transport time is dominated by a small number of slow steps; removal of arginines by mutagenesis eliminates these, revealing a large number of much faster steps.

To summarise so far: arginines, which have a fixed positive charge, have profound effects on the process of transport; that is, in respect of both the rate and the average number of steps required to transport a given polypeptide across the membrane.

### Lysine is deprotonated to facilitate its passage through SecYEG

It has previously been shown that SecYEG is strongly selective against positively charged ions or residues, but more permeable to anions (Dalal 2009,Nouwen 2009,Schiebel 1992). However, the specific effect of arginine has not hitherto been described. To explore this further, we carried out constant velocity steered molecular dynamics (SMD), to explore how easy it is for different side chains to pass through the channel. Here, a 14-residue stretch of polypeptide, based on the PDB structure 5EUL (Li et al., 2016), containing a residue of interest is pulled through the SecY pore (see Methods), and the total force experienced along this pull coordinate is recorded (Fig. 4a). We found that the total force required to move positive charges across the membrane is significantly higher than any other residue type, but with no difference between lysine and arginine (Fig. 4a). The effect is specifically due to the positive charge: uncharged lysine transports just as easily as other uncharged residues (e.g. Lys^0^ vs Leu in Fig. 4a). We quantified this effect using potential of mean force calculations with the coarse-grained Martini force field (Marrink et al., 2007; Monticelli et al., 2008), which predicts a difference in energy barrier of ca. 10 kJ mol^-1^ between charged and uncharged lysine (Fig. 4b).

**Figure 4.**
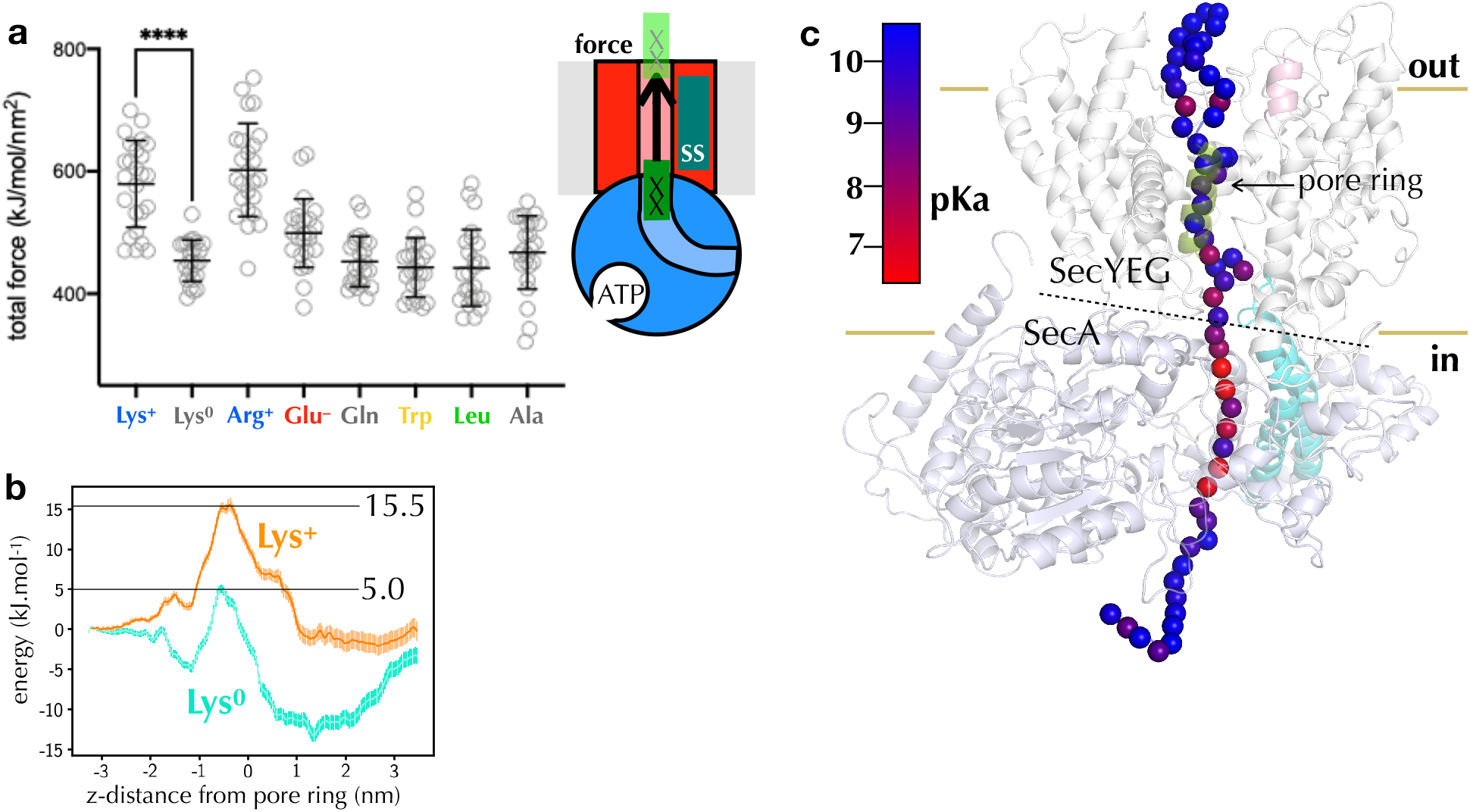
Computational analysis of pre-protein engaged in SecYEG. **a)** Pulling forces for regions of polypeptide containing different residues of interest to pass the SecY pore, as determined using steered MD. Plotted are the integrated forces along the pull coordinated, with 24 repeats for each residue. The mean and standard deviations are shown. **b)** Potential of mean force pathways for a short region of peptide with either protonated (Lys^+^) or deprotonated (Lys^0^) present passing through the SecY pore. These systems were built using the Martini force field. **c)** p*K*_a_ scanning data for lysine residues at different positions along the substrate. Alpha-carbon positions of the substrate are shown as spheres, and coloured according to their calculated p*K*_a_.

The fact that lysine is transported more easily than arginine *in vitro* but not *in silico* would be explained if lysine loses its positive charge before traversing the channel. The p*K*_a_ of lysine in solution is ∼10, but it is readily deprotonated in hydrophobic environments (Isom et al., 2011), whereas the delocalised positive charge of arginine’s guanidinium group (p*K*_a_ ∼ 13.8 (Fitch et al., 2015)) is much harder to remove (Harms et al., 2011). Additionally, the ΔpH component of the PMF would assist with the deprotonation of lysines at the entrance to SecY, and with their rapid reprotonation in the periplasm.

To explore this possibility, we carried out a p*K*_a_ analysis, in which a long stretch of native pre-protein substrate is threaded through the SecA-SecYEG channel (Zimmer et al., 2008), and relaxed over 1 µs of atomistic MD simulation (see Methods). For multiple simulation snapshots, residues along the pre-protein were then mutated to lysine, relaxed with MD, and the p*K*_a_ recorded using the propKa31 program (Søndergaard 2011). The analysis reveals a clear region of p*K*_a_ perturbation at the SecYEG-SecA interface (Fig. 4c), in line with our predictions. Aside from their p*K*_a_s, lysines and arginines are very similar in terms of their physical and chemical properties (see Fig. 2 – figure supplement 2a). The specific deprotonation of lysines therefore seems to be the only plausible explanation for the huge effect arginine relative to lysine on transport time.

### Arginine transport exerts a selection pressure on secreted proteins

If transport of arginines is rate limiting for secretion *in vivo*, one might expect Sec substrates to experience an evolutionary selection pressure to eliminate arginines, where possible – most likely with lysine, the only other positively changed amino acid. To investigate this, we compared the Lys/Arg composition of the mature domain of all known *E. coli* Sec substrates with those that remain in the cytosol. As predicted, secreted proteins have a strong preference for lysine over arginine relative to those in the cytosol (Fig. 5a). We also looked at pre-proteins that are exported, but by the Tat system – i.e independently of Sec – and found their Lys/Arg ratio appears to fall somewhere between Sec substrates and unsecreted proteins (Fig. 5a). This confirms that the Lys/Arg effect is at least in part a product of transport pathway, rather than the extra-cytosolic environment.

**Figure 5.**
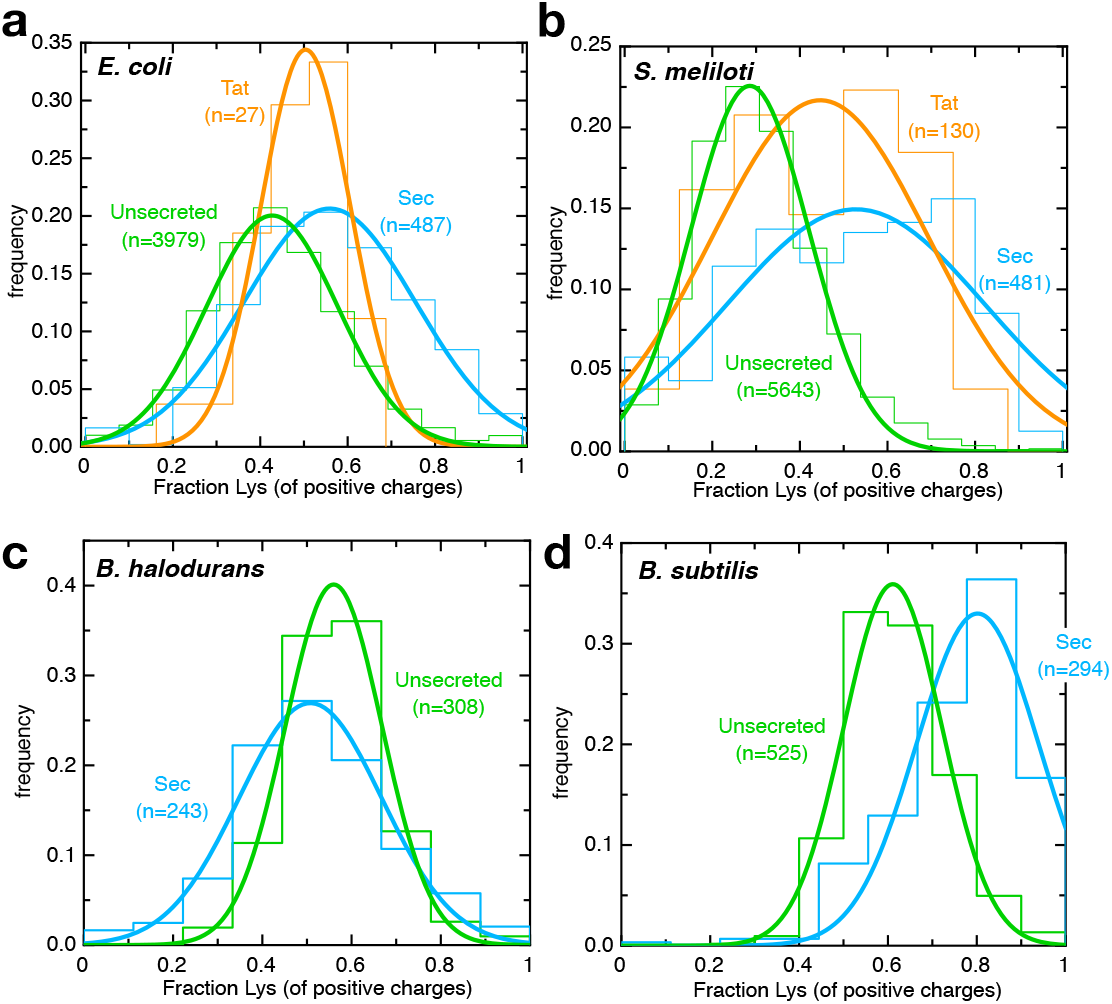
Arginines are selected against for secretion by neutrophiles. **a-d)** Histograms (fine lines) showing fraction of positive residues that are lysine in cytosolic proteins (green), Sec substrates (blue) and Tat substrates (orange; panels **a-b** only), for **(a)** *E. coli*, **(b)** *S. meliloti*, **(c)** *B. halodurans* and **(d)** *B. subtilis*. Best fit single Gaussians are also shown (thick lines) for clarity.

Because *E. coli* only uses Tat to export a very small number of proteins, we carried out the same analysis on a model organism with a large number of annotated Tat substrates (*Sinorhizobium meliloti;* Pickering et al., 2012; Fig. 5b). The results both confirm the *E. coli* observations and show that they hold true across different classes of Gram-negative bacteria. This may be relevant to the mechanism of protein export through Tat, although it could equally reflect differences in the cytosolic vs periplasmic environment.

To explore whether this effect is relates to ΔpH specifically, we carried out the same analysis on an alkalophilic organism with an inverted Δ pH (acid_in_/alkaline_out_), *Bacillus halodurans* (Takami et al., 2000). Consistent with ΔpH being involved in transport, lysine is no longer favoured over arginine for secreted proteins – indeed, the reverse appears to be true (Fig. 5c). Meanwhile, the related bacterium *Bacillus subtilis*, which grows in neutral environments (alkaline_in_/acid_out_), shows the expected increase in lysine for secreted substrates (Fig. 5d). The fact that this trend holds true in three very distantly related bacteria suggests it could be a general feature of secreted proteins.

### Proton motive force speeds up transport primarily through Δψ

The question of how the PMF stimulates SecA-mediated pre-protein transport has been open for decades (Brundage et al., 1990): while it is intuitive to imagine negatively charged residues crossing the membrane electrophoretically with the aid of Δψ, the same effect would equally prevent transport of positive charges. Indeed, a strong electrophoretic pulling force has been observed for negatively charged residues at the SecY channel entrance during co-translational membrane protein insertion, but no equivalent slowing of positively charged residues (Ismail et al., 2015). Deprotonation of lysines at the cytosolic face of SecYEG could neatly circumvent this problem, by imbuing essentially all pre-proteins with a net negative charge as they cross the membrane. To investigate this possibility, we therefore set out to measure the effect of PMF on transport.

To generate a continuous and stable PMF we switched from PLs to IMVs purified from cells (over-)producing SecYEG and 11S (see Pereira et al., 2019). IMVs derived from normally functioning *E. coli* strains contain F_1_F_o_-ATP synthase, which works in reverse to produce a PMF upon addition of exogenous ATP. IMVs differ from PLs in that they contain other, native inner membrane proteins, albeit at much lower stoichiometry relative to SecYEG compared to native membranes. We started by comparing transport of pSpy-pep86 into PLs (Fig. 6a, pink) with transport into IMVs derived from a cell line lacking F_1_F_o_-ATP synthase, and therefore unable to generate PMF from ATP (HB1; green in Fig. 6a). The results show that lag – which corresponds to transport time (see Fig. 2b and Allen et al., 2020) – is identical for both, but IMV transport reaches completion faster, meaning a lower probability that transport stalls or fails. Presumably, this enhanced transport processivity is caused by differences between IMVs and PLs – either a specific effect of auxilliary Sec components, or non-specific differences in the membrane environment.

**Figure 6.**
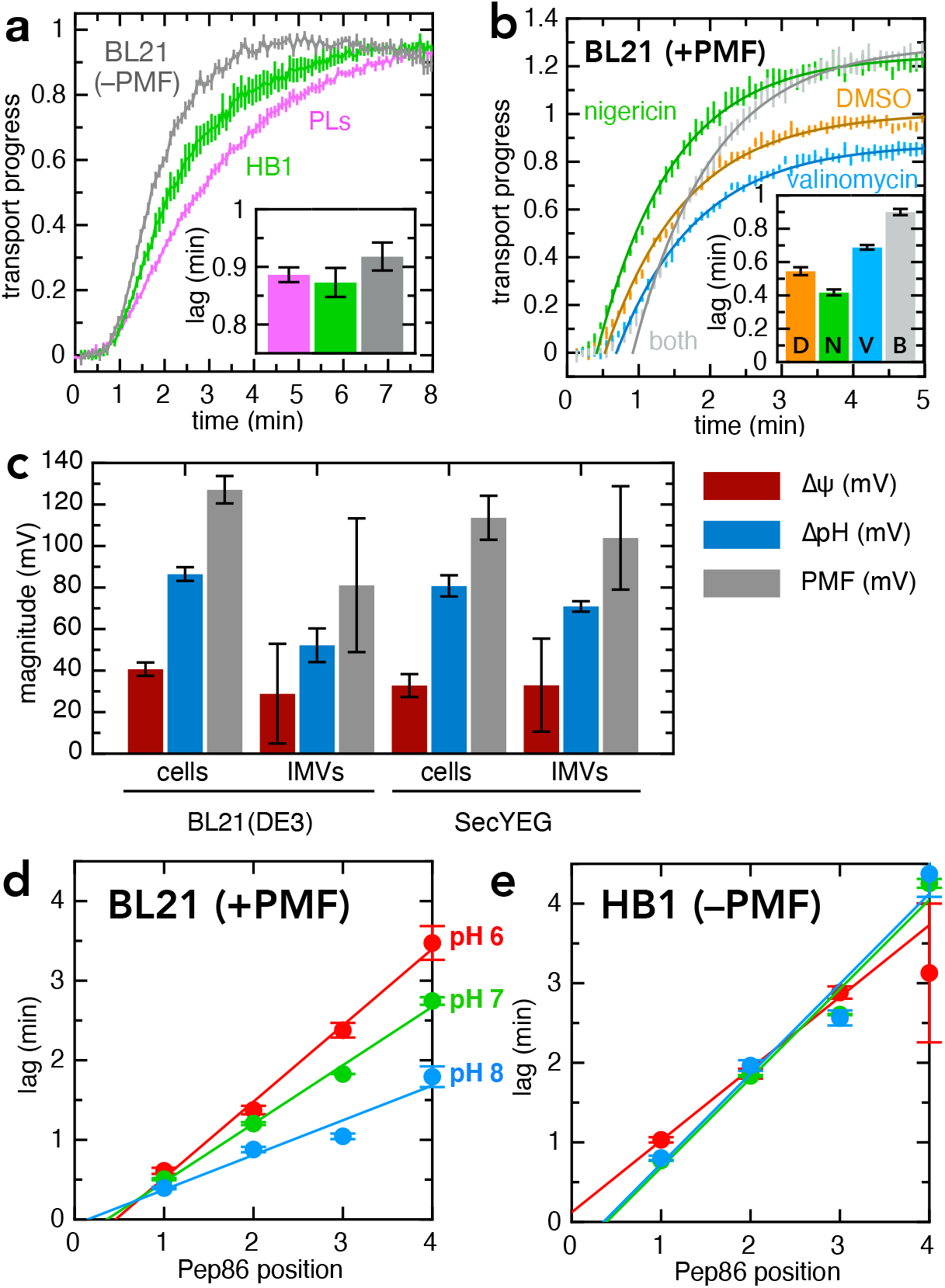
PMF stimulates transport primarily via Δψ. **a)** Transport of pSpy-pep86 into PLs (pink), HB1 IMVs (lacking ATP synthase; green) and BL21 IMVs in the presence of valinomycin and nigericin (grey). Best fit lags (zoomed in for clarity) are shown as an inset. Data points and their heights represent the average and SEM from three (PLs), six (HB1) and six (BL21) experimental replicates. **b)** Transport of pSpy-pep86 into BL21 IMVs in the presence of DMSO (orange), valinomycin (blue), nigericin (green) or both inhibitors (grey). Data points and their heights represent the average and SEM from six replicates, and lines show the best fit to the single exponential + lag model. Best fit lags are shown inset. **c)** Comparison of the proton-motive force (PMF, grey) and its two components (Δψ, red and ΔpH, blue), generated by whole cells and inverted membrane vesicles (IMVs), in BL21(DE3) cells both normal and overexpressing SecYEG. Error bars indicate the standard deviation from three biological replicates (for cells) or three technical replicates (for IMVs). **d)** Lags for import of the pSpy_4x_ series as a function of active pep86 position, into BL21 IMVs at pH 6 (red), 7 (green) and 8 (blue). Data and error bars are the average and SEM from three replicates. **e)** As in panel **c**, but with HB1 IMVs.

In order to compare transport with and without PMF under otherwise identical conditions, we used IMVs with functional F_1_F_o_-ATP synthase (BL21) and either added (–PMF) or omitted (+PMF) two ionophores: valinomycin, a potassium ionophore that specifically depletes Δψ when K^+^ is present in the buffer; and nigericin, an electroneutral K^+^/H^+^ antiporter that dissipates the ΔpH only under the same conditions. Together, these should eliminate PMF entirely; and indeed, as expected, when both inhibitors are present transport into BL21 IMVs give a similar lag to PLs or HB1 IMVs (grey in Fig. 6a, see also inset). In the absence of PMF inhibitors (i.e. +PMF), transport (lag) is about about twice as fast as in their presence (orange vs grey in Fig. 6b). Surprisingly, however, the total amplitude is higher with the inhibitors. As import is largely single turnover under the experimental conditions used (one pre-protein per SecYEG), an increase in amplitude suggests either that pre-proteins are less likely to become irrevocably stuck in the channel during transport, or that more of the SecYEG sites are active.

To separate the effects of Δψ and ΔpH we next measured import with each of the two inhibitors individually. Valinomycin alone (no Δψ, ΔpH increases to compensate) slows transport and produces a small reduction in amplitude (blue in Fig. 6b), while nigericin (no ΔpH, increased Δψ) appears both to speed up transport and increase the maximum amplitude (green in Fig. 6b). These effects are not caused by non-specific effects of the ionophores, as they are not observed for transport into HB1 IMVs (Fig 6, Figure supplement 1a). The faster import with nigericin most likely arises from an increase in Δψ caused by the dissipation of ΔpH; but this cannot explain the increase in amplitude, which is maintained even in the presence of valinomycin. A reasonable hypothesis is that nigericin is helping with the rapid dispersal of local pH fluctuations arising from transport.

While the effects of PMF and its inhibitors are consistent and reproducible, they are relatively small – certainly not sufficient to bridge the difference in rate between ATP-only driven transport *in vitro* and the estimated two orders of magnitude faster transport rate *in vivo* (Allen et al., 2020; Cranford-Smith and Huber, 2018). We therefore measured the magnitude of PMF and its individual components in our purified IMVs, and compared them to the cells from which they are derived (Fig. 6c; see Methods for details). The results suggest that the Δψ produced by the reverse action of ATP synthase in purified IMVs is comparable to that of intact, respiring *E. coli*, and that overexpression of SecYEG also makes little difference (Fig. 6c). However the absolute measured value of Δψ is much lower than is generally reported for E. coli (∼150 mV; McMillan et al., 2007), and it is also possible that protein transport itself consumes PMF *in vitro*, as we have observed for mitochondrial protein import (Ford et al., 2021). Therefore, we cannot conclusively determine the extent to which PMF contributes to pre-protein transport in the absence of auxilliary factors.

### Stimulation of transport by PMF is pH-dependent

To explore the effect of pH further we prepared IMVs at three different pHs (6.0, 7.0 and 8.0) and measured import of the pSpy_4x_ series used previously (Allen et al., 2020) in the same buffer. This series comprises four proteins identical save for the number of copies of mSpy proteins that must pass through SecY before active pep86 becomes available to bind 11S. A plot of lag against active pep86 position therefore gives a straight line with a slope corresponding to the transport rate (Allen et al., 2020).

Just as observed for native pSpy (above), transport is considerably faster (∼2-fold) with PMF (BL21 IMVs) than without (HB1 IMVs) at pH 8.0 (Fig. 6d vs Fig. 6e, blue data). However, as pH decreases, this difference vanishes: at pH 6.0, there is very little PMF effect on rate (Fig 6d-e). The same effect is observed when using ionophores, with the stimulatory effect of nigericin also amplified at high pH (Fig. 6 – Figure supplement 2a-c). Therefore, it seems that the mechanism of transport stimulation by Δψ is pH-dependent. It should be noted that the overall import signal is much lower at low pH (Figure 6 – Figure supplement 2d-e). This could be for any number of reasons, including faster deterioration of SecYEG during the IMV preparation or lower NanoLuc signal at low pH. Once total amplitude is accounted for, however, the rate at which pSpy_4x_ becomes irreversibly trapped in the channel appears to be largely unaffected by pH (Figure 6 – Figure supplement 2f-g).

### PMF stimulation of transport correlates with pre-protein hydrophobicity

We next measured transport of each of the pSpy_XLX_ variants into BL21 IMVs, both in the presence and absence of valinomycin and nigericin, and calculated the stimulatory effect of PMF on transport time for each (see Methods). The results are shown in Fig. 7a-f, with 0% meaning no effect of PMF and 100% a halving of lag in the presence of PMF. As expected, transport is faster in the presence of PMF for all variants. Surprisingly, however, we see little indication that PMF is acting on charged residues. Indeed, all variants where the number of charges is altered show reduced PMF effect relative to native mSpy – even Q→E, which one might reasonably expect to be assisted by Δψ regardless of other factors. Although there are other possible explanations for the lack of PMF stimulation, e.g. differences in folding behaviour that affect how the pre-protein interacts with the Sec system, or a threshold effect that is not reached in our *in vitro* system (see Fig. 6c) these results suggest that the relationship between PMF and charged residues is not straightforward.

**Figure 7.**
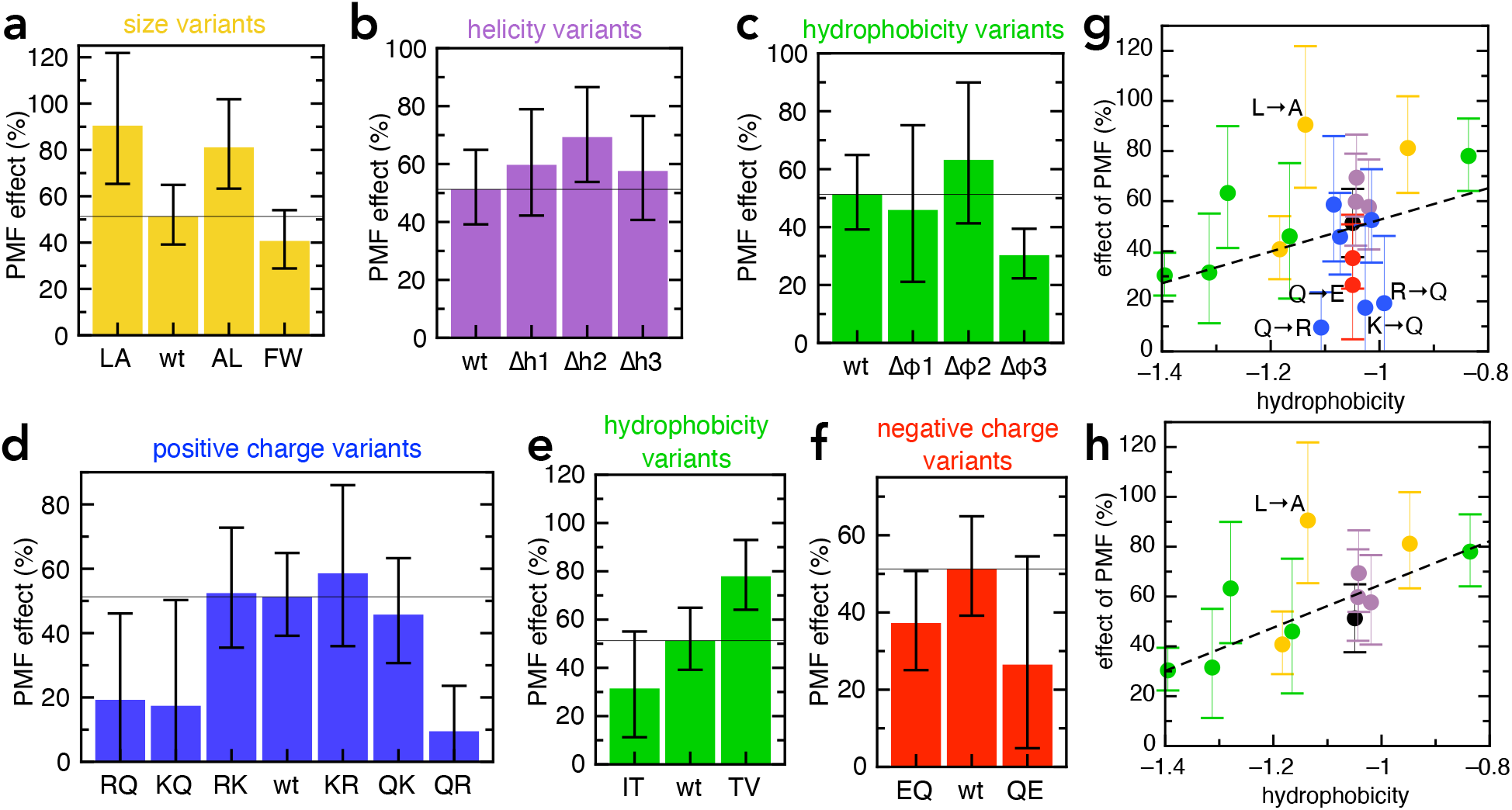
Effect of PMF on transport of the pSpy_XLX_ variants. **a-f)** Stimulatory effect of PMF on transport of the variable region of all the pSpy_XLX_ variants. These were calculated from the difference in lag for import into BL21 IMVs in the presence and absence of valinomycin and nigericin (see Methods for details), where 0% is no difference and 100% is a halving of lag. Error bars are calculated by calculating minimum and maximum values from the SEMs from five replicates of each. **g-h)** PMF effects (coloured as in panels **a-f**) as a function of average hydrophobicity score for the amino acids in each variable mSpy, calculated using the values in (Kyte and Doolittle, 1982). **(g)** Entire data set, with best fit line (r = 0.271). **(h)** Data excluding the charge variants (i.e. the blue and red values; r = 0.681). Outliers are marked directly on the plots.

The only amino acid property that appears to correlate with magnitude of the PMF effect is hydrophobicity, in that more hydrophobic variants exhibit higher stimulation by PMF. A scatter plot of PMF effect vs hydrophobicity shows this more clearly – a straight line through all the points shows a weak correlation (Fig. 7g; r = 0.271), which becomes stronger if the charge variants, which mostly show reduced PMF effect, are omitted (Fig. 7h; r = 0.681). While the large error bars (caused by dividing two experimentally determined numbers) preclude confident assignment of PMF effect to hydrophobic residues, it is the only general amino acid property that has any noticeable effect on this parameter.

## Discussion

The landmark solution of the first atomic structure of SecYEG in complex with SecA, over a decade ago (Zimmer et al., 2008), inspired multiple hypotheses for the underlying mechanism for the transport of pre-proteins across membranes. However, the difficulties obtaining quantitative data on transport itself have, until recently, prevented these from being properly tested. Here, we have applied the newly developed, high time-resolution NanoLuc transport assay (Pereira et al., 2019) to a set of model pre-proteins, designed to capture the rate of transport for amino acid sequences with different properties. Our results reveal a surprisingly simple set of rules for how fast pre-protein is transported through SecYEG: arginine is by far the slowest amino acid to transport, accounting for over 2/3 of the total transport time of the model pre-protein pSpy. Other residues that make a difference are lysines, and bulky residues such as tryptophan, with diffusion of bulky and positive patches probably accounting for the individual steps observed in transport (Fig. 1, purple balls).

In our experiments, transport of arginines is slow and rate-limiting. In contrast lysines behave much more like neutral residues, suggesting that they are at least partially deprotonated before passage across the membrane, a proposition supported by computational analysis of the SecYEG-A structure bound to various model pre-cursors. This deprotonation is presumably facilitated by SecA, with the energy required coming either from ΔpH – protons are more readily removed at the higher pH on the cytosolic side of the membrane – and/or the hydrolytic cycle of ATP, with ATP hydrolysis directly promoting deprotonation. The fact that ΔpH does not appear to have a major effect on transport rate would suggest that the latter is more important, although we interpret these results with caution, given the low magnitude of PMF in our IMV preparations.

In addition to allowing easy passage of lysine through the SecY channel, which is strongly selective against cations, this deprotonation could stimulate transport in two other ways. Firstly, it gives all translocating pre-proteins a net negative charge: as illustrated in Fig. 8, diffusion of negative charges (aspartic and glutamic acid) is likely to be promoted by Δψ (Ismail et al., 2015), while only arginines (which are under-represented in secreted proteins, see Fig. 5) inhibit transport. Additionally, once deprotonated lysines emerge from SecY into the lower pH environment of the periplasm and absent the p*K*_a_ perturbing properties of SecA, they will swiftly be reprotonated and thus unable to diffuse back. This ‘proton ratchet’ effect will prevent backsliding of transported lysines, and thereby contribute to the forward progression of translocation (Fig. 8).

**Figure 8.**
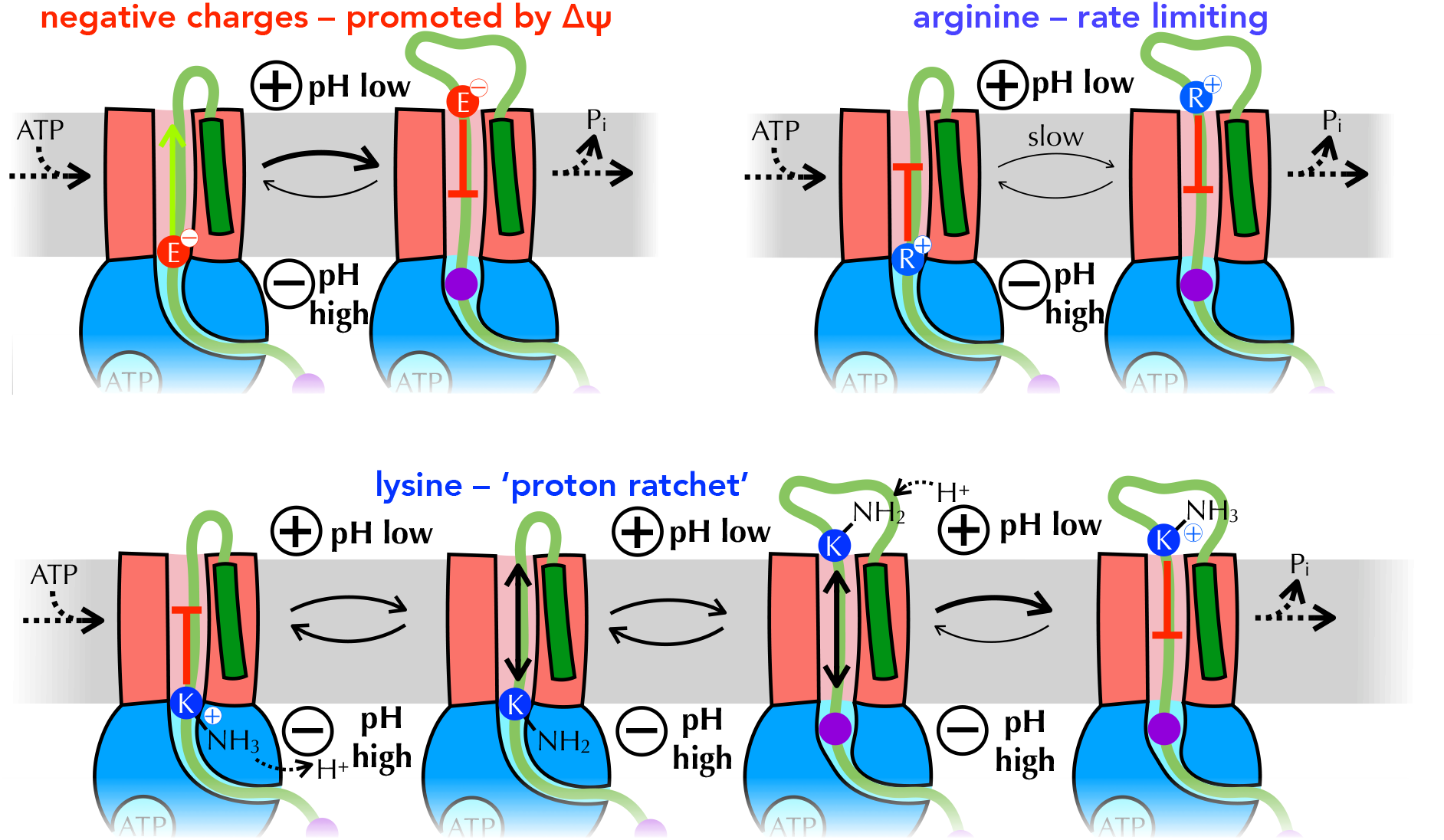
Summary of proposed PMF and ratcheting effects of pre-protein transport. Closeup of step *iii* from Fig. 1, where a negative amino acid (red, top left), arginine (blue, top right) or lysine (blue, bottom) is present at the entrance to SecY. Δψ is expected to promote the forward diffusion of negative charges, and inhibit their reverse movement. Movement of arginine is unfavourable in either direction, with export especially unfavourable when Δψ is present; hence transport of arginine is rate-limiting for the entire process. Lysine is deprotonated at the cytosolic face of SecY (by ΔpH and/or ATP hydrolysis by SecA), allowing its free diffusion across the membrane, and preventing backsliding once it is reprotonated on the other side.

The computational results above were derived from peptides with no other charged residues nearby. Presumably, however, sequence context will have an effect in real life; for example a positive charge adjacent to a negative charge might be harder to deprotonate but easier to transport through the channel, as the two partially cancel each other out. Conversely, a long stretch of consecutive lysines might overwhelm the ability of the Sec machinery to strip and dissipate protons. Poly-lysine is not generally a feature of secreted proteins, but it has previously been used as evidence that positively charged residues are hard to transport (Liang et al., 2012; Nouwen et al., 2009); possibly this strong effect is conferred by the presence of multiple consecutive lysines, as opposed to ones spread evenly through the sequence. The importance of detailed sequence context, along with the compromised PMF in purified IMVs, might also explain the apparently contradictory results we obtain for PMF stimulation of charged pSpy_XLX_ variants.

In addition to the direct mechanistic insights that can be discerned from the above results, they also have more general implications for the nature of protein secretion. It has previously been shown that a lower propensity for folding and the presence of hydrophobic patches are hallmarks of bacterial secreted proteins (Chatzi et al., 2017); to this we can now add a reduction in the number of arginines. Intriguingly, it also seems that any alterations to the charged residues in Spy – either adding or removing positive or negative charges – reduces the magnitude of PMF stimulation. This perhaps indicates that the sequence of Spy is already well optimised for secretion in terms of charge distribution. These observations all imply that ‘secretability’ is an important evolutionary constraint on proteins that localise outside the cytosol, a fact that will have important ramifications for the rational design of secretion-competent proteins for biotechnology or synthetic biology applications. Our results further suggest that replacing arginines with lysines will be a simple but effective way to achieve higher secretability.

Above all, the identification of steps in the translocation process through SecYEG is important because their properties enable us to distinguish between diffusional (described above) and power-stroke (Catipovic 2020,Catipovic 2019) mechanisms of translocation. The steps of the translocation reaction we identify could in theory correspond either to average ratchet lengths during diffusion, or to individual piston motions of a power-stroke. But in the former case the steps would vary with the properties of the translocating pre-protein, as indeed they do; steps in a power stroke, meanwhile, would depend on the geometry of SecA and the conformational changes associated with ATP turnover, and thus be invariant. Ratcheting has the further benefit of allowing other factors to contribute to pre-protein transport, including the PMF; pre-protein folding on the outside over the inside (Corey et al., 2019); and the action of auxiliary Sec components such as SecDF (Tsukazaki, 2018) and periplasmic chaperones (Fürst et al., 2018; Knyazev et al., 2018).

The results presented here might also have interesting implications with respect to the process of membrane protein insertion. While the action of SecA facilitates the transport of positive charges, in its absence – *e*.*g*. during co-translational membrane protein insertion – the process becomes much more problematic. Therefore, the ‘positive-inside rule’ for membrane protein topology (Cymer et al., 2015; vonHeijne, 1989) may in part be a reflection of the fundamental properties of the Sec-machinery.

## Materials and Methods

### Translocation substrate production

To produce the pSpy_XLX_ variants, genes for each variant pSpy were first either synthesised (GeneArt, Thermo Fisher Scientific; pSpy_R→Q_, pSpy_K→Q_, pSpy_R→K_, pSpy_K→R_, pSpy_Q→K_, pSpy_Q→R_, pSpy_E→Q_, pSpy_Q→E_, pSpy_A→L_, pSpy_L→A_, pSpy_I→T_, pSpy_T→V_) or the mutations introduced by site-directed mutagenesis (QuikChange, Agilent; pSpy_Δh1_, pSpy_Δh2_, pSpy_Δh3_, pSpy_Δϕ1_, pSpy_Δϕ2_, pSpy_Δϕ3_, pSpy_F→W_). A gene for pSpy_LX_ was created by removing the dark peptide from pSpy_LD_ (in pBAD/*myc*-His C; Allen et al., 2020) using QuikChange. Each variant mSpy was then cloned between the first mSpy and the pep86 sequences using site directed ligase independent mutagenesis (Chiu et al., 2004). For pSpy_LXX_, the wild type mSpy gene was instead cloned after the pep86 sequence. A complete list of new protein sequences is shown in Supplementary Table 1.

All translocation substrates (including the pSpy_4x_ variants; Allen et al., 2020) were expressed as for native pSpy (Pereira et al., 2019), then purified using a new, streamlined protocol. Cell pellets were resuspended 1:7 (v/v) in 7 M guanidine.HCl on ice, incubated for 30 min, then centrifuged at 100,000 g for 30 mins. The supernatents were next bound to Ni^+2^ affinity resin (∼10x bead volume) by rotating gently for 30 mins at 4 °C, followed by decanting into empty 10 ml gravity flow columns and washing with ≥ 5 bed volumes tris/urea buffer (20 mM Tris-HCl pH 8.0, 6 M urea) supplemented with 30 mM imidazole. Protein was eluted using tris/urea buffer supplemented with 330 mM imizadole, then passed over a Q-sepharose column to remove nucleic acid contamination. Finally, the purified proteins were concentrated and imidazole removed by spin concentration.

### Production of other translocation reagents

Inverted membrane vesicles (IMVs) overexpressing SecYEG and membrane-tethered 11S (∼11S) were produced from previously described cells strains (Pereira et al., 2019). Cells were grown at 37 °C to mid-log phase (OD_600_ ∼ 0.6) in 2xYT supplemented with 100 mg.ml^-1^ amplicillin and 50 mg.ml^-1^ kanamycin, then SecYEG expression was induced with 0.1% (w/v) arabinose. After 2 h, the cells were cooled to 20 °C then ∼11S expression induced overnight (∼16 h) with 1 mM IPTG before harvesting. For purification, a standard protocol was followed (Corey et al., 2018), but using as buffer either M_6_KM (20 mM MES-KOH pH 6.0, 50 mM KCl, 2 mM MgCl_2_), H_7_KM (20 mM HEPES-KOH pH 6.0, 50 mM KCl, 2 mM MgCl_2_) or M_8_KM (20 Mm MOPS-KOH pH 8.0, 50 mM KCl, 2 mM MgCl_2_), depending on the desired final pH.

All other transport reaction components were produced as described previously (Pereira et al., 2019).

### Transport assays

Transport assays were performed and analysed exactly as in (Allen et al., 2020), with 2 µM final concentration of pre-protein (unless otherwise stated) to ensure saturation. For IMV experiments, IMVs and the appropriate pH buffer (M_6_KM, H_7_KM or M_8_KM; pH 8 used unless otherwise stated) were substituted for PLs and TKM (20 mM Tris-HCl pH 8.0, 50 mM KCl, 2 mM MgCl_2_), but all other reaction conditions were kept identical. Where inhibitors were used, they were added from a 100x stock in DMSO to final concentrations of 1 µM (valinomycin) or 2 µM (nigericin), and the corresponding amount of DMSO was added to comparison experiments.

For Berkeley Madonna analysis of the pSpy_XLX_ variants, the model in (Allen et al., 2020) was modified to allow different translocation parameters for the native and variant sequences. A visual representation of this new model is shown in Fig. 3 – Figure Supplement 1b; the complete model is shown in Supplementary Models.

To calculate PMF effect, the import lag for pSpy_LXX_ was subtracted from the lag for each pSpy_XLX_ variant, both in the absence (+PMF) and presence (–PMF) of nigericin and valinomycin, to give transport time just for the variable region. We then calculated PMF effect (in %) as:

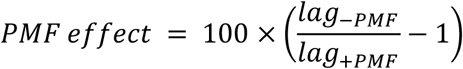

Errors in lag were take as the SEM of lag from four technical replicates, where +PMF and –PMF were identical other than the addition of inhibitors, and error bars for PMF effect were generated by calculating upper and lower bounds from these errors.

### Sequence analysis

Protein properties were calculated over a 9 residue window using ProtScale (https://web.expasy.org/protscale/), using values for hydrophobicity from (Kyte and Doolittle, 1982), helical propensity from (Deléage and Roux, 1987) and bulkiness from (Zimmerman et al., 1968). The same scales were used to plot Figure 2 – Figure supplement 2a.

### Arginine/Lysine ratio determination

The complete proteomes of *E. coli* (strain K12) and *S. meliloti* (strain 1021) were downloaded from UniProt (The UniProt Consortium, 2017) on 1^st^ May 2018 and sorted according whether they are secreted by Sec or Tat system, or unsecreted. For *E. coli*, which is well annotated by UniProt, this was done using information in the ‘Signal Peptide’ column. For *S. meliloti*, secreted substrates were selected according to UniProt ‘Signal Peptide’, then classified as Tat if validated as such in (Pickering et al., 2012), or Sec otherwise. *B. halodurans* and *B. subtilis* sequences were downloaded on 13^th^ March 2019 and analysed as for *E. coli*, omitting signal peptides flagged as ‘Tat-type’. For secretion substrates, the residues corresponding to the SS were removed prior to analysis. The proportion of positive residues that are lysine (Lys/(Lys+Arg)) was calculated for each protein, then plotted as a histogram for each data set (with sample size n): the number of bins was determined by Sturge’s rule (number of bins = 1 + 3.322 * log(n)), and frequency calculated by dividing the population of that bin by n. Fits to single Gaussian curves were performed using Pro Fit (Quansoft)

### Molecular dynamics of substrate moving through SecY

Systems were built using the coordinates of SecYE and a 14 residue stretch of preprotein (residues 778 to 791) from PDB 5eul (Li et al., 2016). The pre-protein was changed into a model peptide, with the sequence AGSGSGSGSGGXGA, where X is the residue of interest (K, R, E, Q, W, L or A). The protein coordinates were built into a POPE:POPG membrane using CHARMM-GUI (Jo et al., 2007; Lee et al., 2016). Proteins were described using the CHARMM36m force field (Best et al., 2012), and waters were TIP3P, with K^+^and Cl^-^ions added to 0.15 M. The protein side chains were set to their default protonation states, as predicted using propKa3 (Søndergaard et al., 2011), apart from the Lys^0^ residue where included. Systems were energy minimized using the steepest descents method, and subsequently equilibrated with 1000 kJ.mol^-1^.nm^-2^ positional restraints on protein backbone atoms for 5 ns, and then relaxed using production MD for 15 ns in the NPT ensemble at 310 K with the V-rescale thermostat and semi-isotropic Parrinello-Rahman pressure coupling. Time steps of 2 fs were used.

For each residue, 24 independent steered MD simulations were run for each pre-protein, where the substituted residue was pulled in a z-axis direction (up through the channel) using an umbrella potential moving at a rate of 1 nm.ns^-1^, with a force constant of 1000 kJ.mol^-1^.nm^-2^. For each repeat, the total pulling force (taken as the area under the curve) was recorded.

### Modelling of an engaged pre-protein in the SecA-SecYEG complex for pK_a_ analysis

Initial coordinates for SecA-SecYEG were taken from chains A, C, D and E of PDB 3DIN (Zimmer et al., 2008), with the ADP-BeF_x_ molecule replaced with ATP (Piggot et al., 2012). Simulations were run over 1 µs in a POPC membrane with explicit waters and Na^+^and Cl^-^ions to 0.15 M using the OPLS-AA force field (Jorgensen et al., 1996), see (Allen et al., 2016) for full details. Taking a 1 µs snapshot as a fully equilibrated starting model, a region of pre-protein 76 residues long was built in an extended configuration through the SecA-SecYEG complex, as described previously (Corey et al., 2016a). The pre-protein was positioned such that it contacted previously identified crosslinking sites in both SecY and SecA (Corey et al., 2016a; Park et al., 2013). The N-terminal SS was modelled as a helix and sited in the SecY lateral gate a per cryo-EM density (Park et al., 2013).

The SecYEG-SecA-pOA-ATP model was then embedded in a POPC membrane, solvated with explicit waters and Na^+^and Cl^-^ions to 0.15 M, and subjected to 1 µs MD simulation. Simulations were carried out as previously described (Allen et al., 2016), using the OPLS-AA force field (Jorgensen et al., 1996), in Gromacs 5.0.4 (Berendsen et al., 1995).

### pK_a_ scanning pipeline

To predict the p*K*_a_ of charged residues as they traverse the SecA-SecYEG complex, we constructed a computational pipeline. For 20 different structural snapshots over the final 500 ns of the SecYEG-SecA-pOA simulations, each of the 76 residues in the pre-protein were substituted to lysine in turn, using Scwrl4 (Krivov et al., 2009), for a total of ca. 1,500 snapshots. These were then relaxed for 1 ns using MD, and the p*K*_a_ of the target lysine determined using propKa31 (Søndergaard et al., 2011).

### Construction of 1D free energy profiles

To provide a more detailed view of the energetic cost of transporting protonated lysine, we constructed free energy profiles of protonated and deprotonated lysine residues through the channel, using the Martini 2.2 force field (Marrink et al., 2007; Monticelli et al., 2008). Following conversion to Martini, elastic bonds of 1000 kJ mol^-1^ were applied between all backbone beads within 1 nm. Electrostatics were described using the reaction field method, with a cut-off of 1.1 nm using a potential shift modifier, and van der Waals interactions were shifted from 0.9-1.2 nm. Simulations were run in the NPT ensemble, with V-rescale temperature coupling at 323 K and semi-isotropic Parrinello-Rahman pressure coupling.

We used steered MD to construct a 1D reaction coordinate for an Ala-Lys-Ala peptide through the SecY channel, with the collective variable constructed from the z-distance between each of the backbone beads in the tripeptide and 5 backbone beads forming the SecY pore (Met 75, Ile 78, Ile 183, Val 278 and Ser 401 in *G. thermodentrifinicans numbering*). We performed 200 ns umbrella sampling MD, using a z-axis umbrella force constant of 2000 kJ mol^-1^ nm^-2^, in 0.1 nm windows along this coordinate, with the lysine either protonated or deprotonated. Construction of the 1D free-energy profile was achieved for the last 150 ns of each window using the weighted histogram analysis method, implemented in the gmx wham program (Hub et al., 2010). Convergence was determined through analysis histogram overlap.

### Proton-motive force measurements

The PMF of *E. coli* whole cells was determined using the distribution of [^14^C]benzoate and [^14^C]methyltriphenylphosphonium^+^ as previously described (Rao et al., 2001). Cultures were grown exactly as described for the induction of SecYEG, to allow for a better comparison between whole cells and the IMVs from which they are isolated, and adjusted to OD_600_ = 1.0 using fresh expression media before measurement. The internal volume of these cells was estimated using the partitioning of ^3^H_2_O and [^14^C]PEG-4000 as previously described (Rao et al., 2001). The PMF of IMVs was measured using [^14^C]methylamine and [^14^C]potassium isothiocyanate as previously described (Reenstra et al., 1980), with the following modifications: instead of flow dialysis, a vacuum manifold (Millipore) fitted with 0.45 μm HA MF membrane filters (Millipore) was used as previously described (Gebhard et al., 2006). Assays were performed at a membrane concentration of 1 [mg protein]/mL in buffer (50 mM Tris-HCl pH 7.0, 5 mM MgCl_2_, 100 mM KCl). 1 mM ATP was added and 2 minutes later cold 2 mL 0.1 M LiCl was used to stop the reaction and membranes were harvested by vacuum filtration, with a further wash of 2 mL 0.1 M LiCl. 250 nCi (4.5 μM and 4.2 μM of [^14^C]methylamine and [^14^C]potassium isothiocyanate respectively) was added per experiment. The interval volume of 1.09 μL [mg protein]^-1^ has been previously calculated (Reenstra et al., 1980). 2 mL scintillation fluid (Amersham) was added to all samples in 4 mL scintillation vials and counted as previously described (Gebhard et al., 2006; Rao et al., 2001). Protein concentration was determined using the BCA assay (Thermo) using bovine serum albumin as a standard.

## Supporting information

Supplementary Models, Tables and Figures

## Acknowledgments

We thank Gail Bartlett for help with the bioinformatics analysis. This work was funded by Wellcome Trust (104632) and the BBSRC (BB/T006889/1; BB/S008349/1; BB/N015126/1; BB/M003604/1; BB/I008675/1). Extended simulations were run on the ARCHER UK National Supercomputing Service (http://www.archer.ac.uk), provided by HECBioSim, the UK High End Computing Consortium for Biomolecular Simulation (hecbiosim.ac.uk), supported by the EPSRC. For the purpose of Open Access, the author has applied a CC BY public copyright licence to any Author Accepted Manuscript version arising from this submission.

