## Supplementary Models, Tables and Figures for "Rate-limiting transport of positive charges through the Sec-machinery is integral to the mechanism of protein transport"

### Supplementary Data

#### Supplementary Models

#### Model 1: Berkeley Madonna model for XLX variants ($n_{var}$ >= 3)

```
METHOD RK4

STARTTIME = 0
STOPTIME = 10
DT = 0.001

{*** Initial parameters ***}

{rate constants (in µM and min, as appropriate)}
k_on = 0.96
k_off = 0.085

k_step_wt = 4.27359
k_init = k_step_wt
k_block_wt = 0.31
k_fail_wt = 0.24189

k_step_var = 5
k_block_var = 0.1
k_fail_var = 0.1

{supplied concentrations (µM)}
Lg = 0.005 {11S concentration (final)}
A0 = 0.004 {starting SecYEG concentration}
B0 = 1 {starting pre-protein concentration}

{additional parameters}
n_wt = 5
n_var = 5
brightness = 471.082 {how much signal is produced per nLuc}

{*** Reaction setup ***}

{initiate concentrations}
b_quad = A0 + B0 + (k_off / k_on)
INIT C = (b_quad - SQRT((b_quad * b_quad) - (4 * A0 * B0))) / 2
INIT A = A0 - C
INIT B = B0 - C
```

```
INIT D[0..n_wt] = 0
INIT E[1..n_var] = 0

{*** Differential equations ***}

d/dt (A) = (C * k_off) - (A * B * k_on) + (ARRAYSUM(D[*]) * k_fail_wt) +
((ARRAYSUM(E[*]) - E[n_var]) * k_fail_var)
d/dt (B) = (C * k_off) - (A * B * k_on) + (ARRAYSUM(D[*]) * k_fail_wt) +
((ARRAYSUM(E[*]) - E[n_var]) * k_fail_var)
d/dt (C) = (A * B * k_on) - (C * k_off) - (C * k_init)
d/dt (D[0]) = (C * k_init) - (D[0] * k_step_wt) - (D[0] * k_block_wt) -
(D[0] * k_fail_wt)
d/dt (D[1..n_wt-1]) = (D[i-1] * k_step_wt) - (D[i] * k_step_wt) - (D[i] *
k_block_wt)  - (D[i] * k_fail_wt)
d/dt (D[n_wt]) = (D[n_wt-1] * k_step_wt) - (D[n_wt] * k_step_var) -
(D[n_wt] * k_block_var)  - (D[n_wt] * k_fail_var)
d/dt (E[1]) = (D[n_wt] * k_step_var) - (E[1] * k_step_var) - (E[1] *
k_block_var)  - (E[1] * k_fail_var)
d/dt (E[2..n_var-1]) = (E[i-1] * k_step_var) - (E[i] * k_step_var) - (E[i]
* k_block_var)  - (E[i] * k_fail_var)
d/dt (E[n_var]) = (E[n_var-1] * k_step_var)

{*** Output ***}

signal = min(E[n_var], Lg) * brightness
```

### Model 2: Berkeley Madonna model for XLX variants ($n_{var} = 2$)

```
METHOD RK4

STARTTIME = 0
STOPTIME = 10
DT = 0.001

{*** Initial parameters ***}

{rate constants (in µM and min, as appropriate)}
k_on = 0.96
k_off = 0.085

k_step_wt = 4.27359
k_init = k_step_wt
k_block_wt = 0.31
k_fail_wt = 0.24189

k_step_var = 5
k_block_var = 0.1
k_fail_var = 0.1

{supplied concentrations (µM)}
Lg = 0.005 {11S concentration (final)}
A0 = 0.004 {starting SecYEG concentration}
B0 = 1 {starting pre-protein concentration}

{additional parameters}
n_wt = 5
n_var = 2
brightness = 471.082 {how much signal is produced per nLuc}
```

```
{*** Reaction setup ***}

{initiate concentrations}
b_quad = A0 + B0 + (k_off / k_on)
INIT C = (b_quad - SQRT((b_quad * b_quad) - (4 * A0 * B0))) / 2
INIT A = A0 - C
INIT B = B0 - C
INIT D[0..n_wt] = 0
INIT E[1..n_var] = 0

{*** Differential equations ***}

d/dt (A) = (C * k_off) - (A * B * k_on) + (ARRAYSUM(D[*]) * k_fail_wt) +
((ARRAYSUM(E[*]) - E[n_var]) * k_fail_var)
d/dt (B) = (C * k_off) - (A * B * k_on) + (ARRAYSUM(D[*]) * k_fail_wt) +
((ARRAYSUM(E[*]) - E[n_var]) * k_fail_var)
d/dt (C) = (A * B * k_on) - (C * k_off) - (C * k_init)
d/dt (D[0]) = (C * k_init) - (D[0] * k_step_wt) - (D[0] * k_block_wt) -
(D[0] * k_fail_wt)
d/dt (D[1..n_wt-1]) = (D[i-1] * k_step_wt) - (D[i] * k_step_wt) - (D[i] *
k_block_wt)  - (D[i] * k_fail_wt)
d/dt (D[n_wt]) = (D[n_wt-1] * k_step_wt) - (D[n_wt] * k_step_var) -
(D[n_wt] * k_block_var)  - (D[n_wt] * k_fail_var)
d/dt (E[1]) = (D[n_wt] * k_step_var) - (E[1] * k_step_var) - (E[1] *
k_block_var)  - (E[1] * k_fail_var)
d/dt (E[2]) = (E[1] * k_step_var)

{*** Output ***}

signal = min(E[2], Lg) * brightness
```

### Supplementary Tables

#### Supplementary table 1: List of pre-protein sequences used.

Differences from the native pSpy sequence are highlighted in bold. pSpy$_{LXX}$ is identical to pSpy$_{XLX}$, but with Pep86 and its linker (yellow and pink) moved in front of the variable pSpy (black)

| Protein name | Sequence* |
|---|---|
| pSpy$_{XLX}$ (wt)† | MRKLTALFVASTLALGAANLAHAADTTTAAPADAKPMMHHKGKFGPHQDMMFKDLNLTDAQKQQIREIMKGQRDQMKRP<br>PLEERRAMHDIIASDTFDKVKAEAQIAKMEEQRKANMLAHMETQNKIYNILTPEQKKQFNANFEKRLTERPAAKGKMPA<br>TAEADTTTAAPADAKPMMHHKGKFGPHQDMMFKDLNLTDAQKQQIREIMKGQRDQMKRPPLEERRAMHDIIASDTFDKV<br>KAEAQIAKMEEQRKANMLAHMETQNKIYNILTPEQKKQFNANFEKRLTERPAAKGKMPATAEGSGVSGWRLFKKISGSG<br>ADTTTAAPADAKPMMHHKGKFGPHQDMMFKDLNLTDAQKQQIREIMKGQRDQMKRPPLEERRAMHDIIASDTFDKVKAE<br>AQIAKMEEQRKANMLAHMETQNKIYNILTPEQKKQFNANFEKRLTERPAAKGKMPATAEGSGENLYFQGHHHHHH |
| pSpy$_{R \to Q}$ | ADTTTAAPADAKPMMHHKGKFGPHQDMMFKDLNLTDAQKQQI**Q**EIMKGQ**Q**DQMK**Q**PPLEE**QQ**AMHDIIASDTFDKVKAE<br>AQIAKMEEQ**Q**KANMLAHMETQNKIYNILTPEQKKQFNANFEK**Q**LTE**Q**PAAKGKMPATAE |
| pSpy$_{K \to Q}$ | ADTTTAAPADAKPMMHH**Q**GKFGPHQDMMF**Q**DLNLTDAQKQQIREIM**Q**GQRDQMKRPPLEERRAMHDIIASDTFD**Q**VKAE<br>AQIA**Q**MEEQRKANMLAHMETQN**Q**IYNILTPEQK**Q**QFNANFEKRLTERPAA**Q**GKMPATAE |
| pSpy$_{R \to K}$ | ADTTTAAPADAKPMMHHKGKFGPHQDMMFKDLNLTDAQKQQI**K**EIMKGQKDQMK**K**PPLEEKKAMHDIIASDTFDKVKAE<br>AQIAKMEEQ**K**KANMLAHMETQNKIYNILTPEQK**K**QFNANFEK**K**LTE**K**PAAKGKMPATAE |
| pSpy$_{K \to R}$ | ADTTTAAPADAKPMMHHKGKFGPHQDMMF**R**DLNLTDAQKQQIREIM**R**GQRDQMKRPPLEERRAMHDIIASDTFD**R**V**R**AE<br>AQIA**R**MEEQRKANMLAHMETQN**R**IYNILTPEQKRQFNANFE**R**RLTERPAAKGKMPATAE |
| pSpy$_{Q \to K}$ | ADTTTAAPADAKPMMHHKGKFGPH**K**DMMFKDLNLTDA**K**KQ**K**IREIMKGQRD**K**MKRPPLEERRAMHDIIASDTFDKVKAE<br>A**K**IAKMEE**K**RKANMLAHMET**K**NKIYNILTPEQKK**K**FNANFEKRLTERPAAKGKMPATAE |
| pSpy$_{Q \to R}$ | ADTTTAAPADAKPMMHHKGKFGPH**R**DMMFKDLNLTDA**R**KQ**R**IREIMKGQRD**R**MKRPPLEERRAMHDIIASDTFDKVKAE<br>A**R**IAKMEE**R**RKANMLAHMET**R**NKIYNILTPEQKK**R**FNANFEKRLTERPAAKGKMPATAE |
| pSpy$_{E \to Q}$ | ADTTTAAPADAKPMMHHKGKFGPHQDMMFKDLNLTDAQKQQIR**Q**IMKGQRDQMKRPPLE**Q**RRAMHDIIASDTFDKVKA**Q**<br>AQIAKME**Q**QRKANMLAHM**Q**TQNKIYNILTP**Q**QKKQFNANF**Q**KRLT**Q**RPAAKGKMPATAE |
| pSpy$_{Q \to E}$ | ADTTTAAPADAKPMMHHKGKFGPH**E**DMMFKDLNLTDA**E**KQ**E**IREIMKGQRD**E**MKRPPLEERRAMHDIIASDTFDKVKAE<br>A**E**IAKMEE**E**RKANMLAHMET**E**NKIYNILTPEQKK**E**FNANFEKRLTERPAAKGKMPATAE |
| pSpy$_{L \to A}$ | ADTTTAAPADAKPMMHHKGKFGPHQDMMFKD**A**N**A**TDAQKQQIREIMKGQRDQMKRPP**A**EERRAMHDIIASDTFDKVKAE<br>AQIAKMEEQRKANM**A**AHMETQNKIYNI**A**TPEQKKQFNANFEKR**A**TERPAAKGKMPATAE |

| | |
|---|---|
| pSpy$_{A \rightarrow L}$ | ADTTTAAPADAKPMMHHKGKFGPHQDMMFKDLNLTD**L**QKQQIREIMKGQRDQMKRPPLEERRAMHDII**L**SDTFDKVKAE<br>**L**QIAKMEEQRK**L**NMLAHMETQNKIYNILTPEQKKQFN**L**NFEKRLTERPA**L**KGKMPAT**L**E |
| pSpy$_{I \rightarrow T}$ | ADTTTAAPADAKPMMHHKGKFGPHQDMMFKDLNLTDAQKQQ**T**RE**T**MKGQRDQMKRPPLEERRAMHD**TT**ASDTFDKVKAE<br>AQ**T**AKMEEQRKANMLAHMETQNK**T**YN**T**LTPEQKKQFNANFEKRLTERPAAKGKMPATAE |
| pSpy$_{T \rightarrow V}$ | ADTTTAAPADAKPMMHHKGKFGPHQDMMFKDLNL**V**DAQKQQIREIMKGQRDQMKRPPLEERRAMHDIIASD**V**FDKVKAE<br>AQIAKMEEQRKANMLAHME**V**QNKIYNIL**V**PEQKKQFNANFEKRL**V**ERPAAKGKMPA**V**AE |
| pSpy$_{\Delta h1}$ | ADTTTAAPADAKPMMHHKGKFGPHQDMMFKDLNLTDAQKQQIREIMKGQRDQMKRPP**IDD**RR**GV**HDIIASDTFDKVKAE<br>AQIAKMEEQRKANMLAHMETQNKIYNILTPEQKKQFNANFEKRLTERPAAKGKMPATAE |
| pSpy$_{\Delta h2}$ | ADTTTAAPADAKPMMHHKGKFGPHQDMMFKDLNLTDAQKQQIREIMKGQRDQMKRPP**IDD**RR**GV**HDIIASDTFDKVK**GD**<br>**V**QIAKMEEQRKANMLAHMETQNKIYNILTPEQKKQFNANFEKRLTERPAAKGKMPATAE |
| pSpy$_{\Delta h3}$ | ADTTTAAPADAKPMMHHKGKFGPHQDMMFKDLNLTDAQKQQIREIMKGQRDQMKRPP**IDD**RR**GV**HDIIASDTFDKVK**GD**<br>**V**QIAKMEEQRKAN**VIG**H**VD**TQNKIYNILTPEQKKQFNANFEKRLTERPAAKGKMPATAE |
| pSpy$_{\Delta \phi 1}$ | ADTTTAAPADAKPMMHHKGKFGPHQDMMFKDLNLTDAQKQQIREIMKGQRDQMKRPPLEERRAMHD**QQ**ASDTFDKVKAE<br>AQIAKMEEQRKANMLAHMETQNKIYNILTPEQKKQFNANFEKRLTERPAAKGKMPATAE |
| pSpy$_{\Delta \phi 2}$ | ADTTTAAPADAKPMMHHKGKFGPHQDMMFKDLNLTDAQKQQIREIMKGQRDQMKRPPLEERRAMHD**QQ**ASDTFDK**N**KAE<br>AQ**Q**AKMEEQRKANMLAHMETQNKIYNILTPEQKKQFNANFEKRLTERPAAKGKMPATAE |
| pSpy$_{\Delta \phi 3}$ | ADTTTAAPADAKPMMHHKGKFGPHQDMMFKDLNLTDAQKQQIREIMKGQRDQMKRPPLEERRAMHD**QQ**ASDTFDK**N**KAE<br>AQ**Q**AKMEEQRKANMLAHMETQNK**Q**YN**Q**LTPEQKKQFNANFEKRLTERPAAKGKMPATAE |
| pSpy$_{F \rightarrow W}$ | ADTTTAAPADAKPMMHHKGK**W**GPHQDMM**W**KDLNLTDAQKQQIREIMKGQRDQMKRPPLEERRAMHDIIASDT**W**DKVKAE<br>AQIAKMEEQRKANMLAHMETQNKIYNILTPEQKKQ**W**NAN**W**EKRLTERPAAKGKMPATAE |

*for XLX variants, only the middle (altered) sequence is shown.

†colour scheme: SS, constant mSpy, variable mSpy, linker, Pep86, TEV site and his tag.

### Supplementary table 2: List of best fit parameters for all pSpy$_{XLX}$ variants

| variant | n | $k_{step,var}$ | $k_{block,var}$ | $k_{fail,var}$ |
|---|---|---|---|---|
| wt | 4 | 4.28498 | 0.30999 | 0 |
| LA | 3 | 3.46711 | 0.31739 | 0.17917 |
| AL | 5 | 2.45927 | 0.16917 | 0.09426 |
| FW | 3 | 1.65508 | 0.35573 | 0 |
| IT | 5 | 8.26094 | 0.27787 | 0.27433 |
| TV | 4 | 2.4821 | 0.22277 | 0 |
| EQ | 4 | 4.53412 | 0.27935 | 0.11069 |
| QE | 4 | 4.49511 | 0.15086 | 0.4502 |

| | | | | |
|---|---|---|---|---|
| RQ | 20 | 61.39963 | 0.31351 | 0.33237 |
| KQ | 4 | 6.01896 | 0.31449 | 0.06284 |
| RK | 5 | 6.10382 | 0.30563 | 0.11119 |
| KR | 4 | 2.45832 | 0.20917 | 0.02191 |
| QK | 4 | 2.4356 | 0.19428 | 0.04172 |
| QR | 4 | 0.66534 | 0.13778 | 8.43E-03 |
| Δh1 | 5 | 5.16757 | 0.29908 | 0.05597 |
| Δh2 | 6 | 4.43953 | 0.17154 | 0.05578 |
| Δh3 | 5 | 3.76111 | 0.18966 | 0.08861 |
| Δϕ1 | 3 | 2.68645 | 0.21479 | 0.1217 |
| Δϕ2 | 4 | 3.70325 | 0.22433 | 0.23137 |
| Δϕ3 | 4 | 4.51002 | 0.22845 | 0.20323 |

### Supplementary Figure Legends

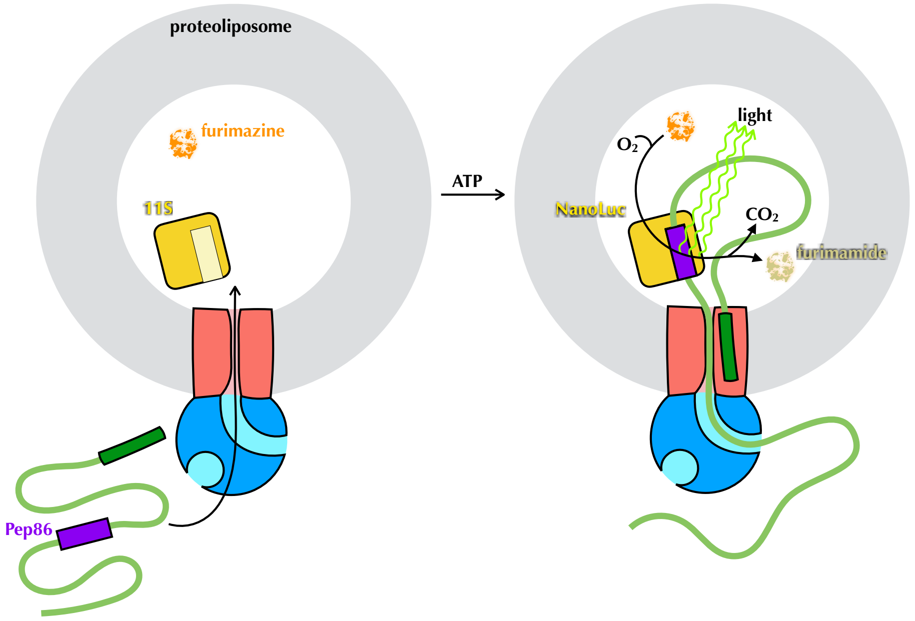


**Figure 1 – Figure supplement 1. The NanoLuc pre-protein transport assay**

Proteoliposomes (grey) with encapsulated 11S (yellow) and SecYEG in the membrane (red) are supplied with SecA (blue), pre-protein (green) containing pep86 (purple) and furimazine (orange). Upon addition of ATP, pre-protein is imported into the PLs, allowing pep86 to complement 11S, poducing NanoLuc. This gives a luminescence signal proportional to the amount of NanoLuc formed, for as long as furimazine and $O_2$ remain in large excess over furimamide and $CO_2$.

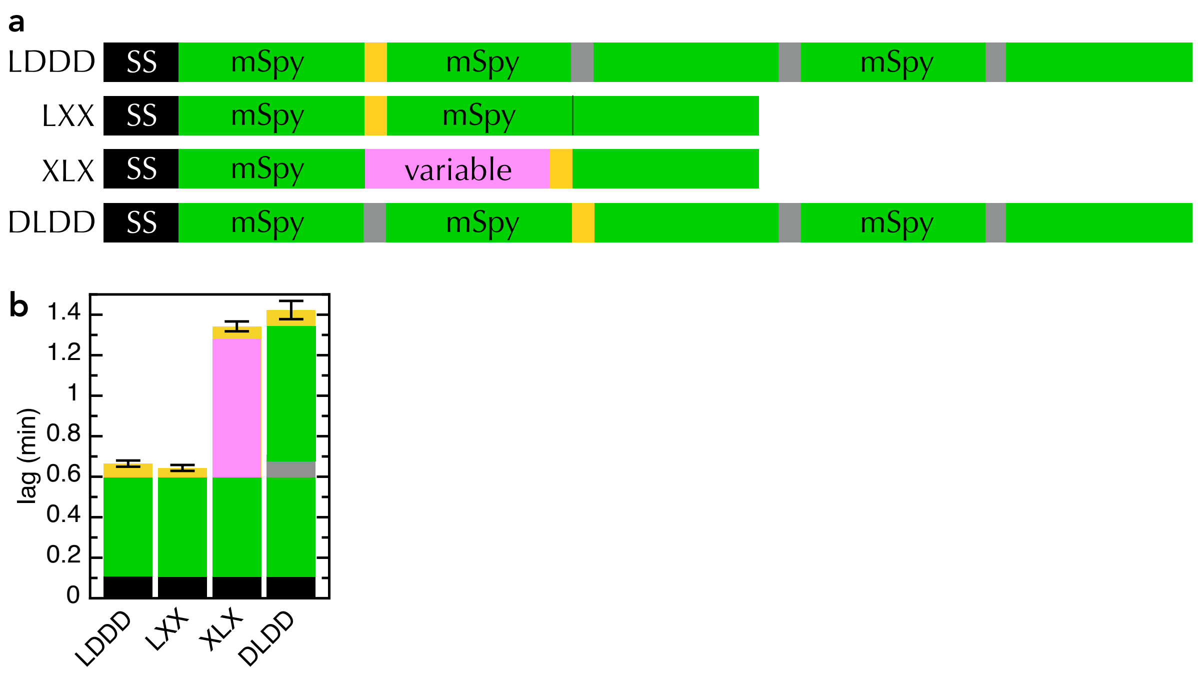


**Figure 2 – Figure Supplement 1. Comparison of new and old transport substrates**

**a)** Schematic of $pSpy_{LXX}$, $pSpy_{XLX}$, $pSpy_{LDDD}$ and $pSpy_{DLDD}$, as in Fig. 1a.

**b)** Transport lags for the pSpy variants in panel **a**. Those for $pSpy_{LXX}$ and $pSpy_{XLX}$ are the same as in Fig. 2c, while those for $pSpy_{LDDD}$ and $pSpy_{DLDD}$ are taken from (Allen et al., 2020).

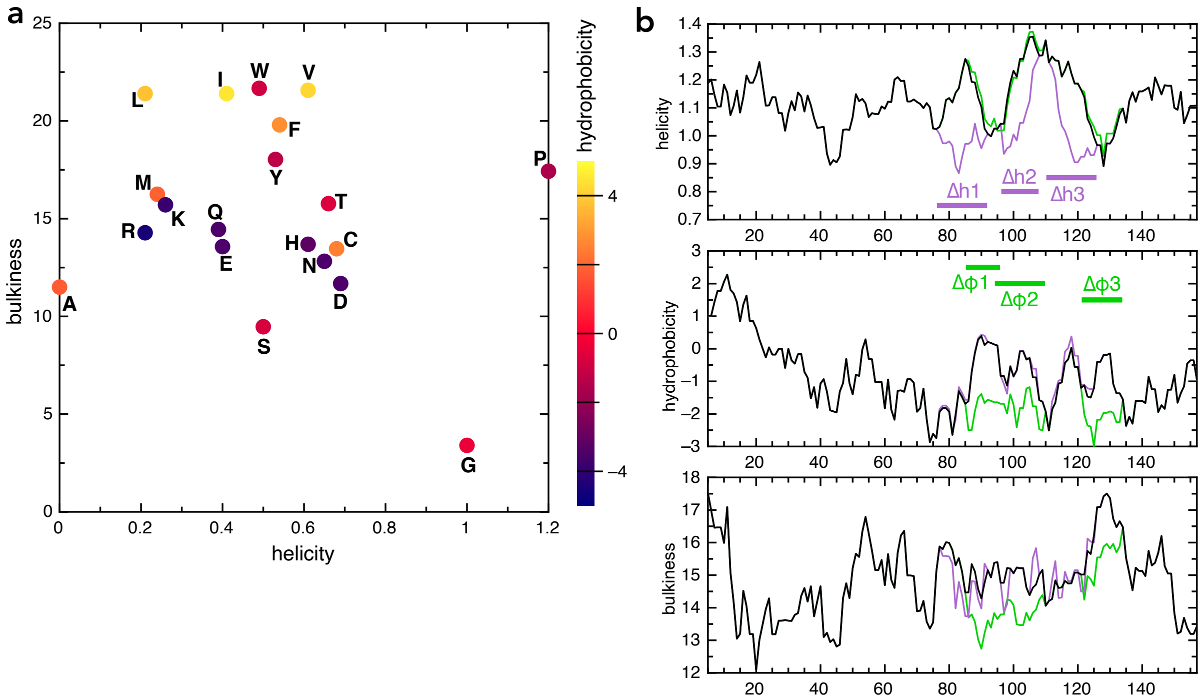


**Figure 2 – Figure Supplement 2. Amino acid and sequence properties**

**a)** Plot of all 20 naturally encoded amino acid by hydrophobicity (scale from Kyte and Doolittle, 1982), helical propensity (scale from Deléage and Roux, 1987) and bulkiness (scale from Zimmerman et al., 1968).

**b)** Plots of helicity (top), hydrophobicity (middle) and bulkiness (bottom) of native pSpy (black) and the helicity (purple) and hydrophobicity (green) variants, as a function of sequence position, averaged over a window of 9 amino acids. Calculations performed using ProtScale on the Expasy portal, using the same scales as in panel **a**.

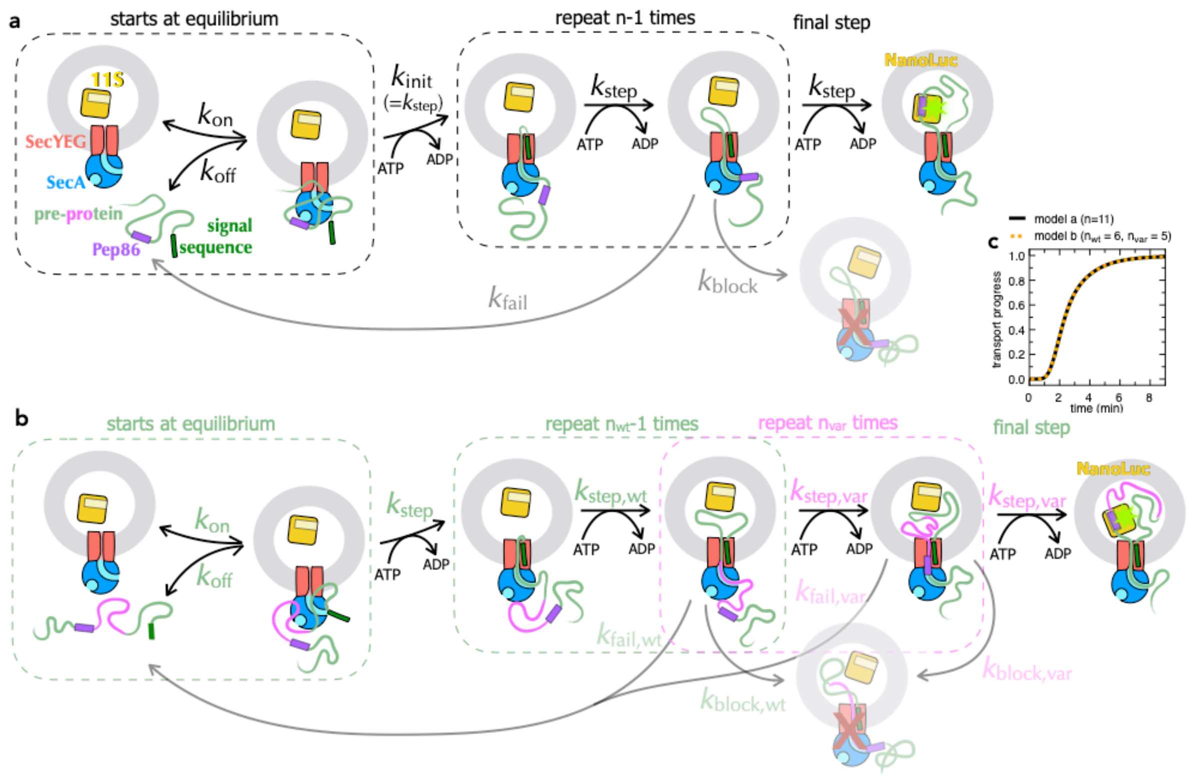


**Figure 3 – Figure Supplement 1. Schematic representations of numerical models**

**a)** Minimal model for transport of pre-protein through the Sec system (Allen et al., 2020). Before the addition of ATP, pre-protein associates with SecYEG and SecA, with an affinity dependent on both the SS and mature domain of the preprotein ($K_d \sim 0.6\ \mu M$ for native proSpy, Allen et al., 2020). Starting the reaction with ATP allows transport to initiate ($k_{init}$, equal to $k_{step}$), followed by a number (n) of steps, each with the rate $k_{step}$. As soon as Pep86 enters the PL it binds to 11S, producing the luminescent signal. Because there are multiple sequential steps before NanoLuc formation, the transport signal begins to appear after a characteristic lag, equal to $k_{init}$ + n × $k_{step}$ (Allen et al., 2020).

Not every transport event that initiates goes on to produce a functional NanoLuc. At any point, transport can fail: either dissociating and requiring reinitiation ($k_{fail}$); or permanently, blocking the channel and preventing further transport ($k_{block}$). In the case of proSpy, each SecYEG appears only to transport a single pre-protein – so the total signal amplitude depends on the number of SecYEG import sites, and the relative rates of transport and $k_{block}$ (Allen et al., 2020). Thus, simple analysis of the NanoLuc traces can provide estimates for the rate of transport and its success rate.

**b)** Extended transport model, separating out transport of the second mSpy of $pSpy_{XLX}$ (pink) from all the other steps (binding, initiation and transport of a single mSpy and Pep86; green). For fitting, all parameters in green are fixed at the values determined from $pSpy_{LXX}$. Brightness is fixed to relative to the wild type $pSpy_{XLX}$ according to the values determined in solution (Fig. 3 – Figure Supplement 3a-f, turquoise bars). The three remaining rate constants ($k_{block,var}$, $k_{step,var}$ and $k_{fail,var}$) are allowed to float, and $n_{var}$ is varied systematically. This model is defined for Berkeley Madonna in Supplementary Models. As long as the wt and var rate constants are all kept the same, model **a** is identical to model **b** if n in **a** is equal to $n_{wt} + n_{var}$ in **b** (see panel **c**).

**c)** Simulated transport curves using the original model (panel **a**) with n = 11 (black solid line), or the split model (panel **b**) with $n_{wt} = 6$ and $n_{var} = 5$ (orange dotted line), and all other parameters kept the same. The fact that these overlay perfectly confirms that the no errors were introduced in extending the model.

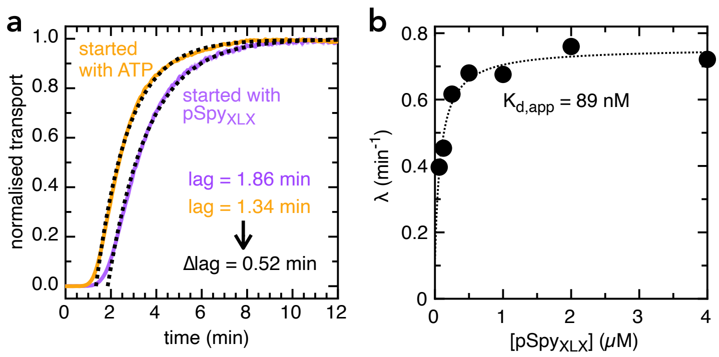


**Figure 3 – Figure Supplement 2. Determination of $k_{on}$ and $k_{off}$ values for $pSpy_{XLX}$**

**a)** Transport of 2 $\mu$M $pSpy_{XLX}$ initiated either by the addition of ATP (orange line) or $pSpy_{XLX}$ (purple line), along with fits (black dotted lines). The difference in lag (0.52 min) corresponds to the time it takes for $pSpy_{XLX}$ to associate with SecYEG-SecA, i.e. $k_{on}^{-1}$ (see Allen et al., 2020 for details). This gives $k_{on}$ = 0.96 $\mu M^{-1}.min^{-1}$, very similar to the value previously determined for native pSpy ($k_{on}$ = 1.25 $\mu M^{-1}.min^{-1}$ Allen et al., 2020): not surprising, given that this interaction is driven primarily by signal sequence and early mature domain which are identical in both proteins.

**b)** λ (the exponential component of the fit, see Allen et al., 2020) as a function of $pSpy_{XLX}$ concentration. Fitting to a simple weak binding equation ($\lambda = \lambda_0 + [pSpy_{XLX}] \div ([pSpy_{XLX}] + K_d)$) gives a good approximation of $K_d$ for the $pSpy_{XLX}$–Sec interaction, as it reflects the starting equilibrium of the binding step (see Fig. 3 – Figure Supplement 1a and Allen et al., 2020). From the best fit value ($K_d$ = 0.089 $\mu$M) and $k_{on}$ (0.96 $\mu M^{-1}.min^{-1}$, panel a), we estimate $k_{off}$ = 0.085 $min^{-1}$ (from the definition $K_d = k_{off} \div k_{on}$).

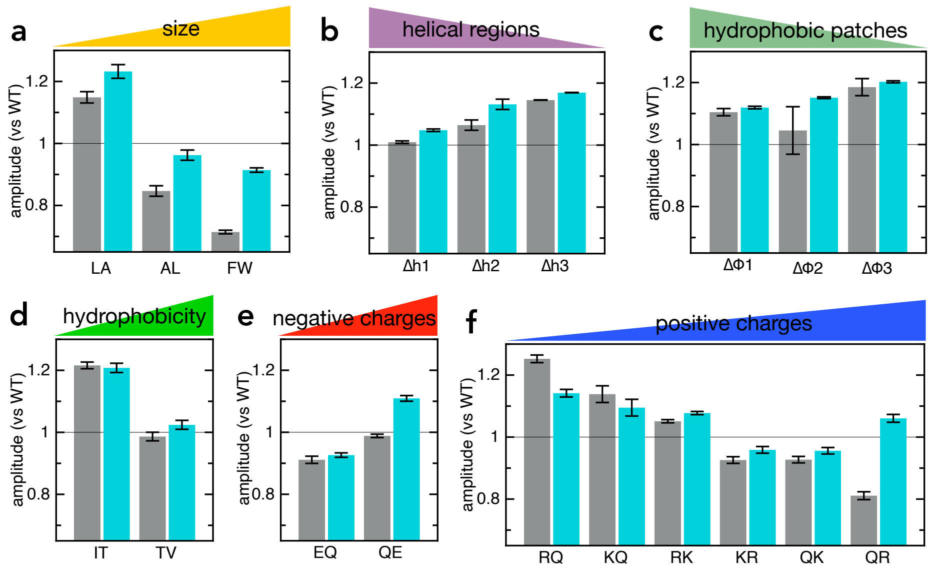


**Figure 3 – Figure Supplement 3. Amplitudes for transport vs binding**

**a-f)** Grey bars: amplitudes for the import of the various $pSpy_{XLX}$ variants (same primary data as in Fig. 2), normalised to the amplitude of native $pSpy_{XLX}$ (measured on the same plate in the same experiment). Turquoise bars: amplitude for binding of 2 $\mu$M of each $pSpy_{XLX}$ variant to 20 pM 11S free in solution. In each case, the signal from 20 pM 11S with only buffer is subtracted, and the amplitude normalised to the value of native $pSpy_{XLX}$ (both measured on the same plate in the same experiment). Error bars are the SEM from four experimental replicates (eight for native $pSpy_{XLX}$ and $pSpy_{LXX}$). With a few exceptions, transport and binding amplitude correspond well, indicating that differences in amplitude are due to differences in the luminescence of NanoLuc (brightness in the model in Figure 3 – Figure Supplement 1).

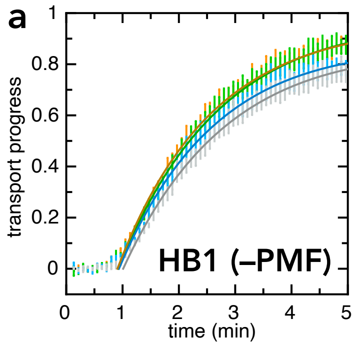


**Figure 6 – Figure Supplement 1. Transport into HB1 IMVs with ionophores**

Exactly as in Fig 6b, but with HB1 IMVs instead of BL21 IMVs. In this case, inhibitors have little or no effect on the kinetics of import.

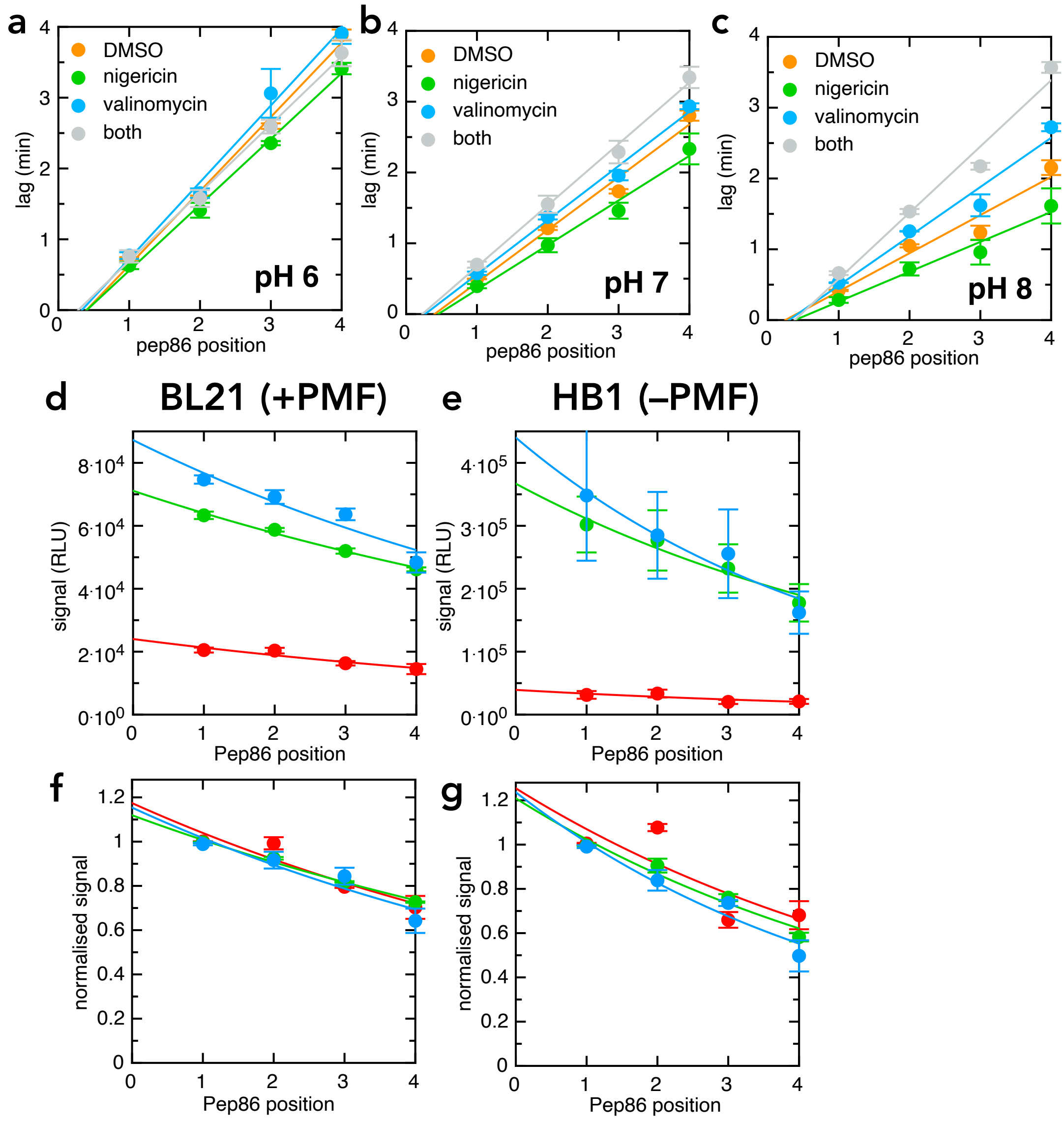


**Figure 6 – Figure Supplement 2. Additional LDDD series transport traces.**

**a-c)** Transport lag for the LDDD series in the presence of DMSO (orange), nigericin (green), valinomycin (blue) or nigericin and valinomycin together (grey), at pH 6 (panel **a**), pH 7 (panel **b**) or pH 8 (panel **c**). Data points and error bars are the average and SEM from three experimental replicates (six for DMSO).

**d-g)** Amplitudes from the same primary experiments for which lag was determined in Figs 6c-d. **(d-e)** are unnormalised, and show that pH strongly reduces total signal both in BL21 **(d)** and HB1 **(e)** IMVs. **(f-g)** are normalised to the value for $pSpy_{LDDD}$, and show that the probabilities of pre-protein becoming trapped in the channel (reflecting $k_{block}$) are indistinguishable at all three pH values.

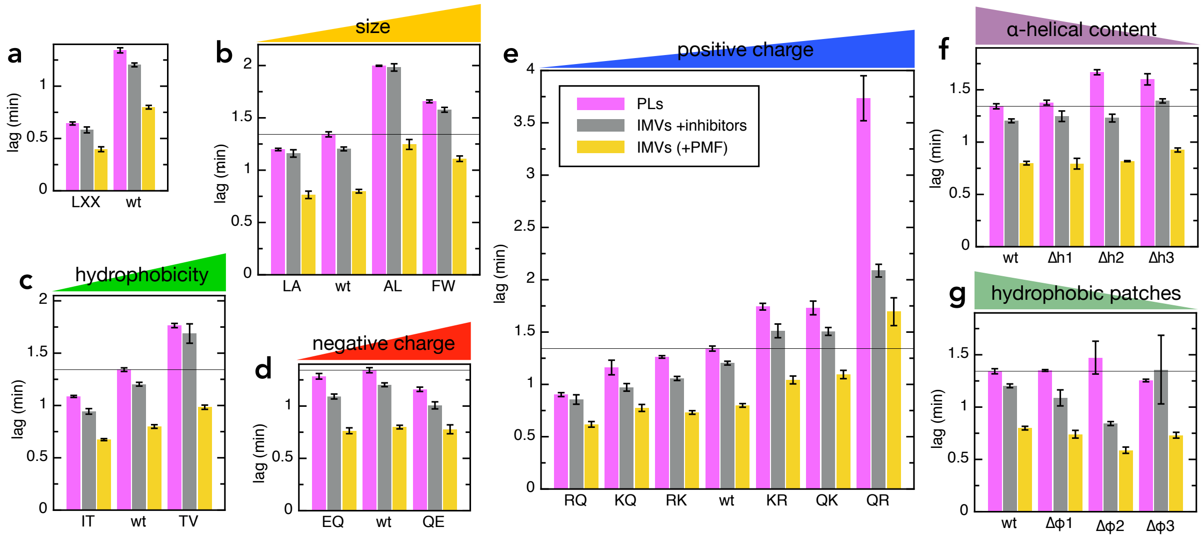


**Figure 7 – Figure Supplement 1. Comparisons of PLs and BL21 IMVs ± ionophores**

**a-g)** Lags for transport of all pSpy$_{XLX}$ variants into PLs (pink; same data as in Fig. 2) and BL21 IMVs in the absence (yellow) or presence (grey) of valinomycin and nigericin (raw data for the PMF stimulation data in Fig. 7).
